# Dual and Opposing Roles for the Kinesin-2 Motor, KIF17, in Hedgehog-dependent Cerebellar Development

**DOI:** 10.1101/2022.08.06.502363

**Authors:** Bridget Waas, Brandon S. Carpenter, Olivia Q. Merchant, Kristen J. Verhey, Benjamin L. Allen

## Abstract

While the kinesin-2 motors KIF3A and KIF3B have essential roles in ciliogenesis and Hedgehog (HH) signal transduction, potential role(s) for another kinesin-2 motor, KIF17, in HH signaling have yet to be explored. Here, we investigated the contribution of KIF17 to HH-dependent cerebellar development, where *Kif17* is expressed in both HH-producing Purkinje cells and HH-responding cerebellar granule neuron progenitors (CGNPs). Germline *Kif17* deletion in mice results in cerebellar hypoplasia due to reduced CGNP proliferation, a consequence of decreased HH pathway activity mediated through decreased Sonic HH (SHH) protein. Notably, Purkinje cell-specific *Kif17* deletion phenocopies *Kif17* germline mutants. Surprisingly, CGNP-specific *Kif17* deletion results in the opposite phenotype– increased CGNP proliferation and HH target gene expression due to altered GLI transcription factor processing. Together these data identify KIF17 as a key regulator of HH-dependent cerebellar development, with dual and opposing roles in HH-producing Purkinje cells and HH-responding CGNPS.

**Teaser:** During cerebellar development, the KIF17 microtubule motor performs opposing roles in HH-producing and HH-responding cells.

## Introduction

Hedgehog (HH) signaling is a major mitogenic stimulus for postnatal expansion of the developing cerebellum (*1, 2*). Sonic hedgehog (SHH) ligand is produced by Purkinje cells and promotes cerebellar granule neural progenitor (CGNP) proliferation (*1-3*). *Shh* deletion within Purkinje cells results in cerebellar hypoplasia and reduced CGNP proliferation (*3*). Conversely, increasing the dosage of *Shh* in Purkinje cells results in cerebellar hyperplasia, as well as the formation of additional cerebellar lobes (*4*). Genetic deletion of other HH pathway components, namely *Gli2* (a key transcriptional effector of the HH pathway), *Gas1* or *Boc* (essential HH pathway co-receptors), within CGNPs leads to cerebellar hypoplasia due to reduced CGNP proliferation (*5, 6*).

In addition to CGNPs, mature cerebellar granule neurons (CGNs) and Bergmann glia (BG) are HH-responsive (*6*). Recent work demonstrated that abrogating HH signaling within BG (through conditional *Smo* deletion) results in a non-cell autonomous reduction in CGNP proliferation and mild patterning abnormalities (*7*). Notably, the role of HH signaling within mature CGNs remains unclear.

A key organelle that is required for proper HH signaling in mice is the primary cilium (reviewed in (*8*)). Primary cilia are microtubule-based organelles that project from the cell surface and act as signaling centers for the HH pathway. Anterograde transport within primary cilia is accomplished by the heterodimeric kinesin-2 motor, KIF3A/KIF3B. Loss of either subunit in mice, *Kif3a* or *Kif3b*, lead to an absence of primary cilia, dysregulation of HH signaling and embryonic lethality (*9-11*). Within the developing cerebellum, loss of *Kif3a* within CGNPs leads to cerebellar hypoplasia due to reduced CGNP proliferation and loss of mitogenic response to SHH ligand (*12*). In addition to KIF3A/KIF3B function in ciliogenesis, KIF3A and its adaptor protein, KAP3, regulate HH signaling by binding to and regulating GLI transcription factors (*13*).

The kinesin-2 motor family contains two additional motor complexes in mammals, heterodimeric KIF3A/KIF3C and homodimeric KIF17 (reviewed in (*14, 15*)). These motors are known as accessory motors, as they do not have clear roles within mammalian ciliogenesis (*16-18*). Loss of *Kif17* is well-tolerated across several model organisms, though KIF17 does have defined roles within several neuronal tissues. Within *Caenorhabditis elegans*, loss of OSM-3, a KIF17 homologue, leads to disruption of the distal region of primary cilia in sensory neurons (*19, 20*). In *Danio rerio*, loss of *Kif17* results in disrupted photoreceptor outer segment development (*21-23*), as well as morphological changes to olfactory cilia (*24*). *Kif17* deletion in mice leads to short term memory issues, learning disabilities and disruption of NR2B trafficking in the hippocampus (*17, 25*). Given that KIF17 can alter primary cilia with functional consequences in multiple neuronal cell types across different species, we investigated the contribution of KIF17 to HH signaling during postnatal cerebellar development.

Here we find that *Kif17* is expressed within SHH-producing Purkinje cells and HH-responsive CGNPs. Germline *Kif17* deletion leads to cerebellar hypoplasia, reduced CGNP proliferation and decreased HH target gene expression across multiple HH-responsive cell types. Purkinje cell-specific *Kif17* deletion phenocopies the germline mutant, demonstrating a requirement for KIF17 in Purkinje cells for proper HH signaling, a finding that correlates with reduced SHH protein levels within Purkinje cells in *Kif17* mutant animals. Conversely, CGNP-specific *Kif17* deletion results in upregulation of HH target genes and increased CGNP proliferation *in vitro* and *in vivo*, a finding that correlates with reduced GLI3 protein levels (a transcriptional repressor of HH signaling). Together these data suggest that KIF17 plays dual roles in HH-dependent cerebellar development– promoting HH signaling in Purkinje cells through the regulation of SHH ligand and restricting HH signaling in CGNPs through the regulation of GLI transcription factor processing.

## Results

### *Kif17* is expressed within Purkinje cells and cerebellar granule neural progenitors and is required for normal cerebellar development

To investigate a role for the kinesin-2 motor, KIF17, in HH signal transduction, we generated *Kif17* mutant mice on a congenic C5BL/6J background. Similar to previous work on *Kif17* (*17, 22*), but in contrast to genetic deletion of other kinesin-2 family members (*9, 10*), *Kif17* homozygous mutant animals are viable and fertile, with no gross morphological abnormalities. Expression analysis revealed that *Kif17* is expressed within the developing cerebellum, starting at postnatal day 4 (P4) and continuing into adulthood (Supplemental Figure 1A-F). X-GAL staining of *Kif17*^*+/+*^ and *Kif17*^*lacZ/lacZ*^ pups at P10 demonstrated *Kif17* expression within the Purkinje cell layer (PCL) and to a lesser degree within the external granule layer (EGL; Figure 1A-B). *Kif17* is expressed in a graded fashion along the anterior-posterior axis, with the strongest signal detected within the posterior lobes (Figure 1A-B), similar to what has been reported for the HH pathway target *Gli1* (*6*). To evaluate if the loss of KIF17 impacted HH-dependent cerebellar development, we continued our analysis of *Kif17*^*-/-*^ cerebella at postnatal day 10, following the peak of HH-dependent CGNP proliferation.

**Figure 1.**
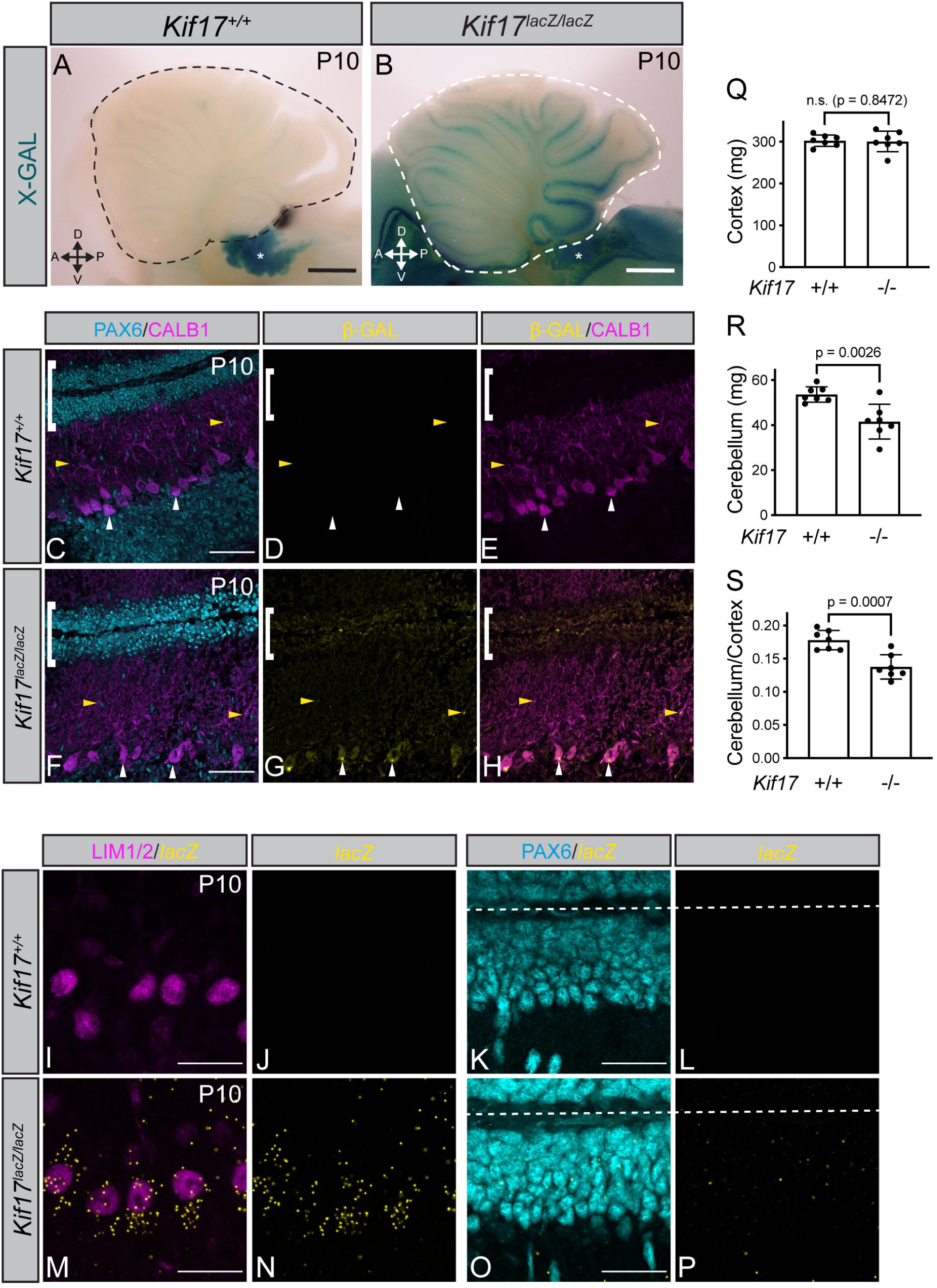
*Kif17* is expressed within Purkinje cells and cerebellar granule neural progenitors and is required for normal cerebellar development. Whole-mount X-gal staining of *Kif17*^*+/+*^ (**A**) and *Kif17*^*lacZ/lacZ*^ (**B**) cerebella at postnatal day 10 (P10). Scale bar, 500 μm. Asterisks denote endogenous Beta galactosidase (β-GAL) activity in the choroid plexus (*64*). Immunofluorescent antibody detection of β-GAL (yellow) in *Kif17*^*+/+*^ (**C-E**) and *Kif17*^*lacZ/lacZ*^ (**F-H**) P10 posterior cerebellar lobes. Antibody detection of PAX6 (cyan) and Calbindin (CALB1, magenta) mark granule neuron nuclei and Purkinje cells, respectively. White brackets denote the external granule layers (EGL); white arrowheads indicate Purkinje cell bodies and yellow arrowheads indicate Purkinje cell dendrites. Scale bars (**C, F**), 50 μm. Fluorescent *in situ* detection of *lacZ* mRNA (yellow; I-P) in *Kif17*^*+/+*^ (**I-L**) and *Kif17*^*lacZ/lacZ*^ (**M-P**) P10 cerebella. Antibody detection of LIM1/2 (magenta; **I, M**) and PAX6 (cyan; **K, O**) identify Purkinje cell and cerebellar granule neural progenitor nuclei, respectively. Dashed lines separate individual external granule layers. Scale bars (**I, K, M, O**), 25 μm. Quantitation of cortex weight (**Q**), cerebellar weight (**R**), and cerebellar weight normalized to cortex weight (**S**) in P10 *Kif17*^*+/+*^ and *Kif17*^*-/-*^ mice. Data are mean ± s.d. Each dot represents an individual animal. *P*-values were determined by a two-tailed Student’s *t*-test.

To identify which cell(s) express *Kif17*, we performed immunofluorescence for beta-galactosidase (β-GAL) in *Kif17*^*+/+*^ and *Kif17*^*lacZ/lacZ*^ cerebella at P10 (Figure 1C-H). In posterior lobes of *Kif17*^*lacZ/lacZ*^ cerebella, we observed punctate localization of β-GAL within the cell bodies of Purkinje cells and in a subset of dendrites (Figure 1G, arrowheads). Further, we observed β-GAL signal within cerebellar granule neuron progenitors (CGNPs) of the EGL (Figure 1G, bracket). To confirm expression within these two cell populations, we performed fluorescence *in situ* hybridization in posterior lobes of *Kif17*^*+/+*^ and *Kif17*^*lacZ/lacZ*^ cerebella (Figure 1I-P). In *Kif17*^*lacZ/lacZ*^ cerebella, we detected *lacZ* punctae surrounding Purkinje cell nuclei (Figure 1M-N) and CGNP nuclei (Figure 1O-P), corroborating the B-GAL localization results. Notably, *Kif17* expression persists in Purkinje cells through P21 (Supplemental Figure 1G-L). Finally, RT-qPCR analysis confirmed *Kif17* expression in CGNPs and verified efficient *Kif17* deletion in mutant animals (Supplemental Figure 1M-O). Together, these data indicate that *Kif17* is expressed in two cell populations in the developing cerebellum– SHH-producing Purkinje cells and SHH-responsive CGNPs.

Analysis of cortical (Figure 1Q) and cerebellar (Figure 1R) weights at P10 indicated that *Kif17* mutant cerebella are significantly smaller than *Kif17*^*+/+*^ littermates. Notably, this difference persisted even after normalizing cerebellar weight to cortical weight (Figure 1S). No significant difference in cortices or cerebellar weights were detected in *Kif17*^*+/-*^ animals (Supplemental Figure 1P-R). Notably, cerebellar hypoplasia was still observed in *Kif17* mutant animals maintained on a mixed C57BL/6J; 129S4/SvJaeJ background (Supplemental Figure 1S). However, this phenotype was not observed in *Kif17* mutants maintained on a congenic 129S4/SvJaeJ background (Supplemental Figure 1T). Together, these data suggest that KIF17 promotes cerebellar development, albeit in a genetic background-dependent fashion.

### *Kif17* germline deletion results in reduced CGNP proliferation and decreased *Gli1* expression within all HH-responsive cells

To further investigate which layers of the cerebellum are affected by KIF17 loss, we measured the thickness of the Purkinje cell layer (PCL, Figure 2A, Supplemental Figure 2A), which is composed of Purkinje cells and Bergmann glia. Additionally, external granule layer thickness was measured (EGL, Figure 2B, Supplemental Figure 2B) where CGNPs reside. Although we did not detect significant changes in PCL thickness, we did observe statistically significant reductions in EGL thickness within both posterior and anterior lobes of *Kif17*^*-/-*^ cerebella (Figure 2B, Supplemental Figure 2B). Since previous work demonstrated that reduced EGL thickness is associated with a reduction in CGNP proliferation (*5*), we next examined *in vivo* proliferation of CGNPs in *Kif17*^*+/+*^ and *Kif17*^*-/-*^ P10 cerebella. Within the posterior lobes, we observed a significant reduction in the percentage of Ki67^+^ cells and EdU^+^ cells (Figure 2C-J). Within the anterior lobes, we similarly observed a significant reduction in CGNP proliferation (Supplemental Figure 2C-D), although to a lesser degree. Intriguingly, while there is a significant reduction in the percentage of EdU^+^ cells, we also observed decreased EdU fluorescence within posterior and anterior lobes of *Kif17*^*-/-*^ cerebella (Supplemental Figure 2E-F). Altogether, these data suggest that cerebellar hypoplasia in *Kif17*^*-/-*^ mice is due to reduced CGNP proliferation.

**Figure 2.**
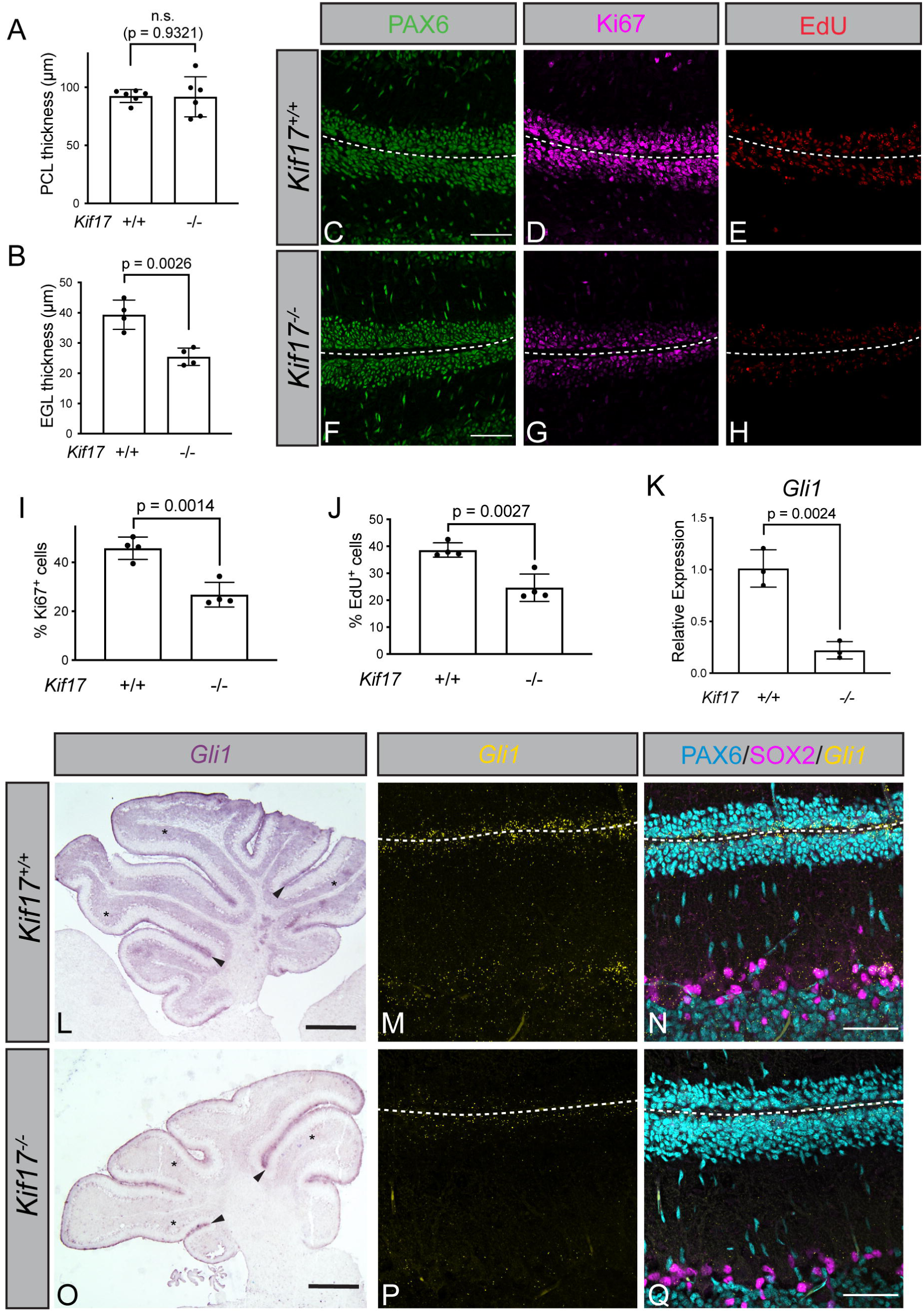
*Kif17* germline deletion results in reduced CGNP proliferation and decreased *Gli1* expression within all HH-responsive cells. Quantitation of Purkinje cell layer (PCL, **A**) and external granule layer (EGL, **B**) thickness in posterior lobes of P10 *Kif17*^*+/+*^ and *Kif17*^*-/-*^ cerebella. Immunofluorescent analysis of CGNP proliferation in the posterior lobes of P10 *Kif17*^*+/+*^ (**C-E**) and *Kif17*^*-/-*^ (**F-H**) cerebella. Antibody detection of PAX6 (green; **C, F**) and Ki67 (magenta; **D, G**). Fluorescent azide detection of EdU (red; **E, H**). Scale bars (**C, F**), 50μm. Dashed line separates individual external granule layers. Percentage of Ki67^+^ (**I**) and EdU^+^ (**J**) cells within the posterior lobes of P10 *Kif17*^*+/+*^ and *Kif17*^*-/-*^ cerebella. RT-qPCR detection of *Gli1* expression (**K**) in P10 *Kif17*^*+/+*^ and *Kif17*^*-/-*^ cerebella. Data are mean ± s.d. Each dot represents the average of 3-5 images per individual animal (**A-B, I-J**) or an individual animal (**K**). *P*-values were determined by a two-tailed Student’s *t*-test. (**L, O**) *In situ* hybridization detection of *Gli1* in *Kif17*^*+/+*^ (**L**) and *Kif17*^*-/-*^ (**O**) P10 cerebella. Arrowheads point to EGL (CGNPs), while asterisk denotes inner granule layer (CGNs). Scale bars (**L, O**), 500 μm. Fluorescent *in situ* detection of *Gli1* (yellow; **M-N, P-Q**) and antibody detection (**N, Q**) of PAX6 (cyan) and SOX2 (magenta) to label granule neurons and Bergmann glia, respectively. Scale bars (**N, Q**), 50 μm. Dashed lines separate external granule layers.

To determine whether decreased CGNP proliferation was associated with alterations in the levels of HH signaling in *Kif17*^*-/-*^ cerebella, we quantified expression of the HH target gene, *Gli1*, using RT-qPCR and found it is significantly decreased in *Kif17*^*-/-*^ P10 cerebella (Figure 2K). Expression of other HH target genes, *Ptch1, Ptch2, Ccnd1*, also trend lower in *Kif17*^*-/-*^ cerebella (Supplemental Figure 2G-I). Since *Gli1* is expressed in several HH-responsive cells in the developing cerebellum [CGNPs, Bergmann glia and cerebellar granule neurons (CGNs)], section *in situ* hybridization for *Gli1* was performed to define which cell population(s) displayed downregulated *Gli1* expression. Surprisingly, we found that *Gli1* expression is reduced across all HH-responsive cells (Figure 2L-Q). Additionally, reduced *Gli1* expression persists in CGNs and Bergmann glia in P21 *Kif17*^*-/-*^ cerebella (Supplemental Figure 2J-M). Since we did not observe *Kif17* expression within Bergmann glia and CGNs, we hypothesized that KIF17 acts in a non-cell autonomous fashion in SHH-producing Purkinje cells to regulate *Gli1* expression.

### Purkinje cell-specific *Kif17* deletion results in a non-cell autonomous HH loss-of-function phenotype

To directly assess KIF17 function in Purkinje cells, we conditionally deleted *Kif17* within Purkinje cells using a *Shh*^*Cre*^ driver (Figure 3A). The specificity of *Shh*^*Cre*^ was confirmed through breeding with *Rosa26*^*LSL-tdT*^ reporter mice (Supplemental Figure 3A-F). Consistent with previous reports (*26*), *Shh*^*Cre*^ efficiently mediates recombination in Calbindin (CALB1)-positive Purkinje cells. Importantly, *Shh*^*Cre*^ is a loss-of-function allele; however, reducing *Shh* dosage does not alter cerebellar size in *Kif17*^*-/-*^*;Shh*^*+/-*^ pups compared to *Kif17*^*-/-*^ littermates (Supplemental Figure 3G). RT-qPCR analysis revealed significantly reduced *Kif17* expression in *Shh*^*Cre*^*;Kif17*^*fl/fl*^ cerebella (Figure 3B), suggesting efficient deletion within Purkinje cells (note that residual *Kif17* expression is likely due to the presence of *Kif17*-expressing CGNPs). Remarkably, Purkinje cell-specific *Kif17* deletion results in cerebellar hypoplasia, phenocopying *Kif17* germline deletion (Figure 3C, Supplemental Figure 3H, cf. Figure 1S).

**Figure 3.**
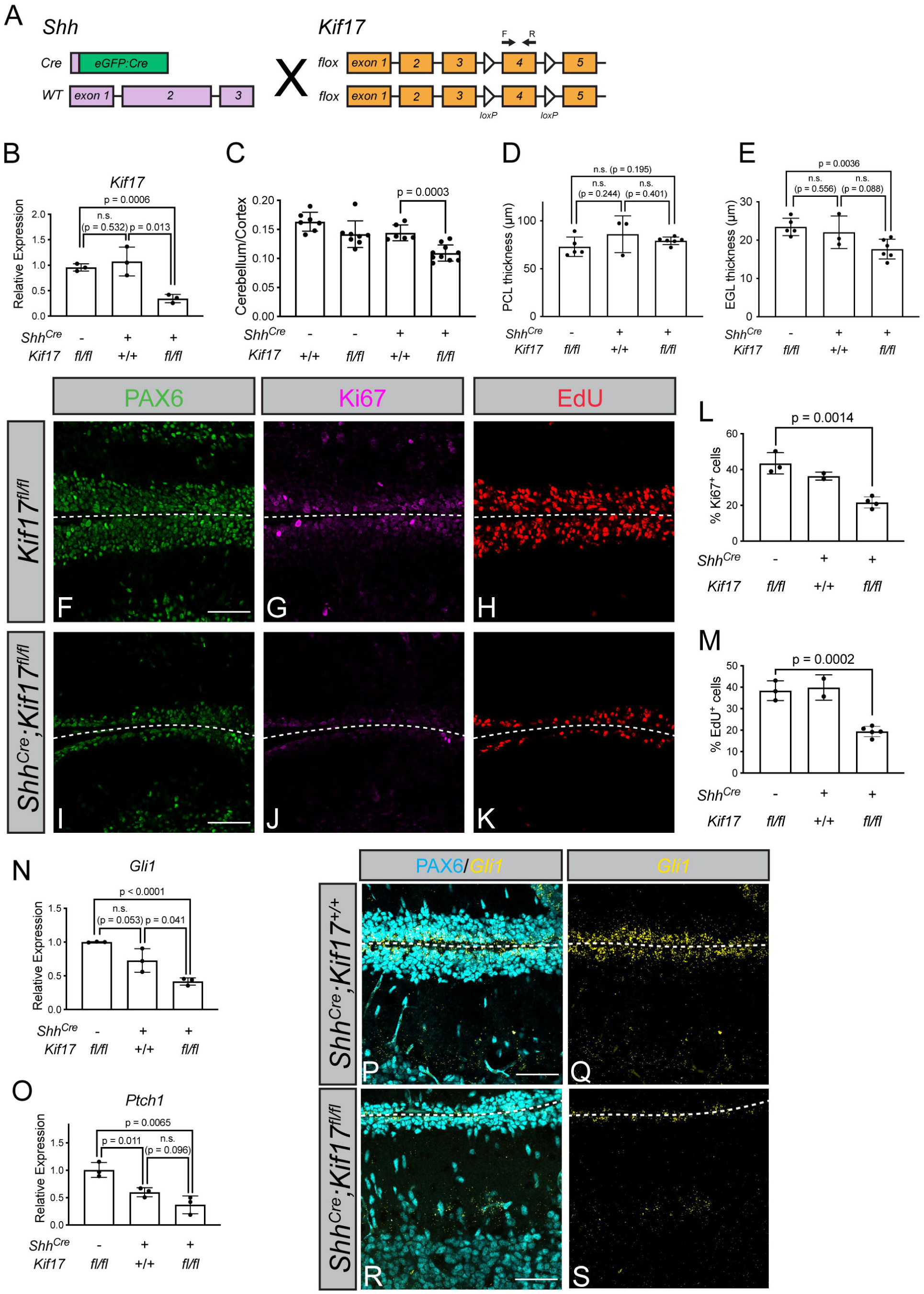
Purkinje cell-specific *Kif17* deletion results in a non-cell autonomous HH loss-of-function phenotype. (**A**) Schematic representing conditional *Kif17* deletion within Purkinje cells using *Shh*^*Cre*^. Arrows above exon 4 denote qPCR primers. Relative *Kif17* expression (**B**) by RT-qPCR in *Kif17*^*fl/fl*^, *Shh*^*Cre*^*;Kif17*^*+/+*^ and *Shh*^*Cre*^*;Kif17*^*fl/fl*^ P10 whole cerebella. Cerebellum weight normalized to cortex weight (**C**) P10 in P10 control and Purkinje cell-specific *Kif17* deletion mice. Quantitation of PCL (**D**) and EGL (**E**) thickness within posterior lobes of P10 cerebella in *Kif17*^*fl/fl*^, *Shh*^*Cre*^*;Kif17*^*+/+*^ and *Shh*^*Cre*^*;Kif17*^*fl/fl*^ mice. Immunofluorescent analysis of cerebellar granule neural progenitor proliferation in *Kif17*^*fl/fl*^ (**F-H**) and *Shh*^*Cre*^*;Kif17*^*fl/fl*^ (**I-K**) P10 cerebella. Antibody detection of PAX6 (green; **F, I**) and Ki67 (magenta; **G, J**). Fluorescent azide detection of EdU (red; **H, K**). Scale bars (**F, I**), 50 μm. Percentage of Ki67^+^ (**L**) and EdU^+^ (**M**) cells within the posterior lobes in P10 control and conditional *Kif17* deletion cerebella. Relative expression of *Gli1* (**N**) and *Ptch1* (**O**) measured by RT-qPCR in P10 whole cerebella in *Kif17*^*fl/fl*^, *Shh*^*Cre*^*;Kif17*^*+/+*^ and *Shh*^*Cre*^*;Kif17*^*fl/fl*^ mice. Data are mean ± s.d. Each dot represents an individual animal (**B-C, N-O**) or an average of 5 images per animal (**D-E, L-M**). *P*-values were determined by a two-tailed Student’s *t*-test. Fluorescent *in situ* detection of *Gli1* (yellow; **P-S**) and antibody detection of PAX6 (cyan; **P, R**) within posterior cerebellar lobes of P10 *Shh*^*Cre*^*;Kif17*^*+/+*^ (**P, Q**) and *Shh*^*Cre*^*;Kif17*^*fl/fl*^ (**R, S**) mice. Scale bars (**P, R**), 50 μm. Dashed lines separate external granule layers.

As with *Kif17* germline mutants, PCL thickness is unaltered in Purkinje cell-specific *Kif17* mutant pups in either the posterior (Figure 3D) or anterior (Supplemental Figure 3I) lobes. However, there is a significant reduction in EGL thickness, specifically in posterior lobes (Figure 3E, Supplemental Figure 3J). Consistent with *Kif17*^*-/-*^ mice, analysis of CGNP proliferation revealed a significant reduction in the percentage Ki67^+^ cells and EdU^+^ cells within both the posterior and anterior lobes of *Shh*^*Cre:GFP*^*;Kif17*^*fl/fl*^ mice compared to control littermates (Figure 3F-M, Supplemental Figure 3K-L). Additionally, we observed significant reductions in the expression of multiple HH target genes, including *Gli1* and *Ptch1* (Figure 3N-O) as well as *Ptch2* and *Ccnd1*, as measured by RT-qPCR (Supplemental Figure 3M-N). Fluorescence *in situ* hybridization revealed reduced *Gli1* expression in *Shh*^*Cre:GFP*^*;Kif17*^*fl/fl*^ cerebella within CGNPs, BG and CGNs (Figure 3P-S). These data demonstrate that Purkinje cell-specific *Kif17* deletion phenocopies germline *Kif17* mutant cerebella, establishing an essential role for KIF17 within SHH-producing Purkinje cells during cerebellar development.

### KIF17 regulates SHH protein in the developing cerebellum

The reduction of HH target gene expression across multiple HH-responsive cells in *Kif17* mutant cerebella suggested a non-cell autonomous role for KIF17 in HH signal transduction. Given that SHH, the only HH ligand expressed in the developing cerebellum, is produced by Purkinje cells, we explored a role for KIF17 in Purkinje cell regulation of SHH localization and release. Initially, examination of *Shh* expression by RT-qPCR revealed that *Shh* transcripts are downregulated in both *Kif17*^*-/-*^ mice (Figure 4A) and Purkinje cell-specific conditional *Kif17* mutants (Figure 4B). Next, we assessed the protein levels of SHH ligand and observed levels of N-terminal SHH are subtly but not significantly decreased in the cerebella of *Kif17* germline mutants [Figure 4C-D; SHH antibody specificity was validated in *Shh*^*-/-*^ tissue (Supplemental Figure 4A)].

**Figure 4.**
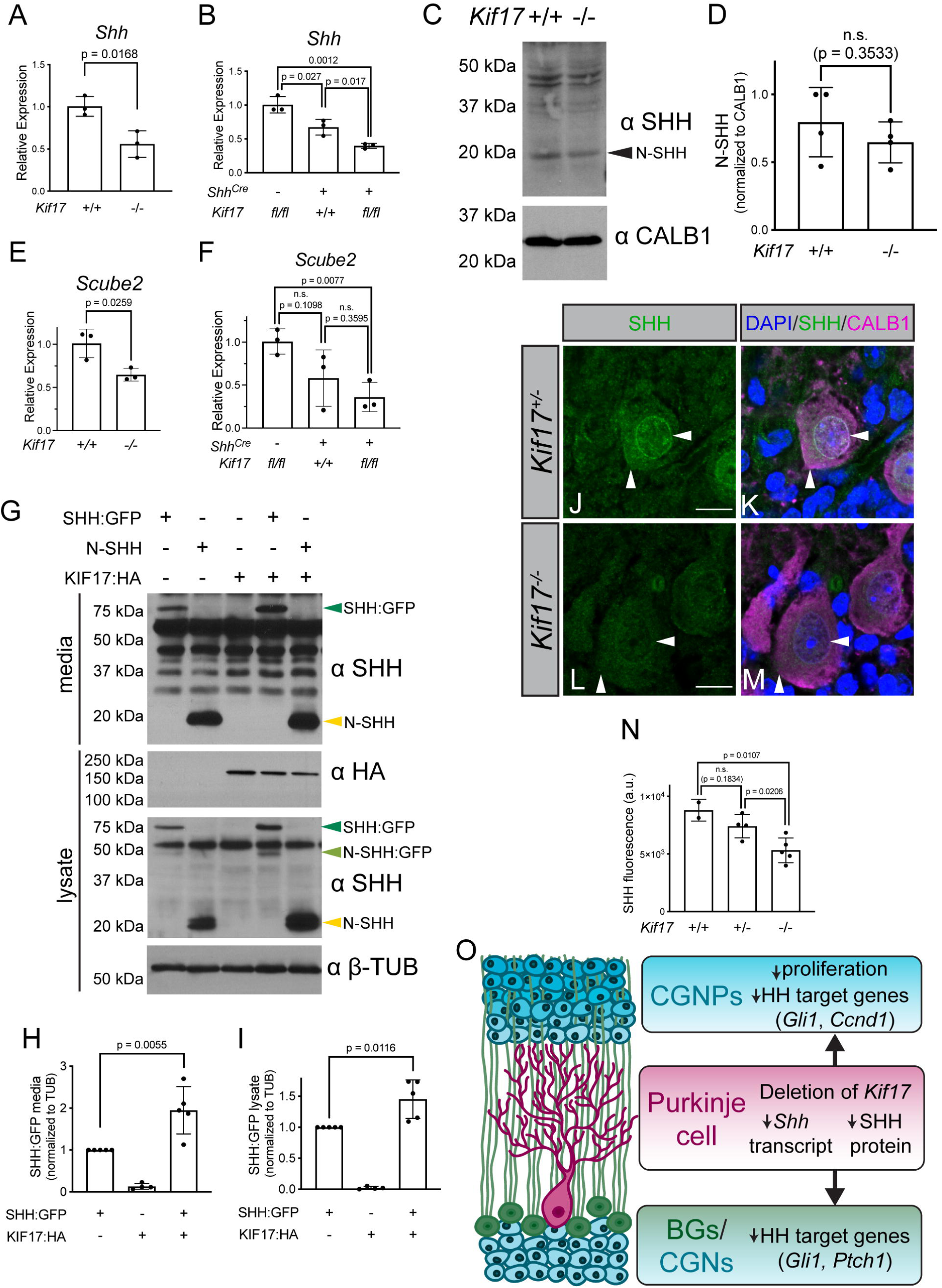
KIF17 regulates SHH protein in the developing cerebellum. Relative expression of *Shh* (**A, B**) by RT-qPCR in P10 whole cerebella of (**A**) *Kif17*^*+/+*^ and *Kif17*^*-/-*^ mice and (**B**) *Kif17*^*fl/fl*^, *Shh*^*Cre*^*;Kif17*^*+/+*^ and *Shh*^*Cre*^*;Kif17*^*fl/fl*^ mice. Western blot analysis examining levels of SHH, using an antibody targeted to the N-terminus of SHH (**C**) in *Kif17*^*+/+*^ and *Kif17*^*-/-*^ P10 cerebella. Antibody detection of Calbindin (⍰-CALB1) was used to confirm equal loading across lanes. Arrowhead denotes secreted N-SHH (19kDa). The molecular masses (in kDa) of protein standards are indicated at the left of each blot. Quantitation of the levels of N-SHH (**D**) normalized to Calbindin in *Kif17*^*+/+*^ and *Kif17*^*-/-*^ in P10 cerebella. RT-qPCR analysis of *Scube2* expression (**E, F**) in P10 whole cerebella of (**E**) *Kif17*^*+/+*^ and *Kif17*^*-/-*^ mice and (**F**) *Kif17*^*fl/fl*^, *Shh*^*Cre*^*;Kif17*^*+/+*^ and *Shh*^*Cre*^*;Kif17*^*fl/fl*^ P10 whole cerebella. Western blot analysis (**G**) of media and cell lysates collected from COS-7 cells expressing HA-tagged KIF17 (KIF17:HA), full length SHH fused to GFP (SHH:GFP) or N-SHH. Blots were incubated with antibodies directed against SHH (⍰-SHH) and HA (⍰-HA). Antibody detection of β-tubulin (⍰ β-TUB) was used to confirm equal loading across lanes. Arrowheads denote full length SHH:GFP (68 kDa), N-SHH:GFP (42 kDa) or N-SHH (19 kDa). The molecular masses (in kDa) of protein standards are indicated at the left of each blot. Quantitation of full length SHH:GFP in the media (**H**) and in the cell lysates (**I**) normalized to β-tubulin within COS-7 cells. Immunofluorescent detection of SHH using an antibody targeted to the C-terminus of SHH (green; **J-M**). DAPI denotes nuclei (blue, **K, M**). Antibody detection of Calbindin, (CALB1, magenta; **K, M**) in P10 posterior cerebellar sections from *Kif17*^*+/-*^ (**J, K**) and *Kif17*^*-/-*^ (**L, M**) mice. Horizontal arrowheads indicate SHH localization to Golgi/ER, while vertical arrowheads denote cytoplasmic localization. Scale bars (**J, L**), 10 μm. Quantitation of SHH fluorescence (**N**) in posterior cerebellar lobes of P10 *Kif17*^*+/+*^, *Kif17*^*+/-*^ and *Kif17*^*-/-*^ mice. Data are mean ± s.d. Each dot represents an individual animal (**A-B, D-F**), independent experiment (**H-I**) or the average of 5 images per animal (**N**). *P*-values were determined by a two-tailed Student’s *t*-test. (**O**) Summary of Purkinje cell-specific *Kif17* deletion on HH ligand production and HH response in the developing cerebellum.

We also examined levels of the HH co-receptor, BOC, which is expressed in Purkinje cells (*5*) and has been recently demonstrated to regulate SHH localization in cytonemes of NIH/3T3 cells (*27*). Notably, levels of *Boc* transcripts (Supplemental Figure 4B-C) and BOC protein are unaltered in *Kif17* deletion cerebella (Supplemental Figure 4D-E). However, *Scube2*, which encodes a key regulator of SHH protein release (*28, 29*), is significantly reduced in P10 cerebella from both *Kif17*^*-/-*^ (Figure 4E) and Purkinje cell-specific *Kif17* mutant animals (Figure 4F). Given the reduced levels of *Scube2*, we speculated that KIF17 could impact SHH ligand release or secretion.

To assess a role for KIF17 in SHH release, we utilized a gain-of-function approach, where COS-7 cells were driven to express epitope-tagged KIF17 (KIF17:HA) and either full-length (SHH:GFP) or N-terminal SHH (N-SHH; Figure 4G). While KIF17 expression does not alter the levels of secreted N-SHH (Supplemental Figure 4F), we did observe increased levels of secreted full-length SHH (Figure 4H). We also observed significantly increased levels of intracellular SHH, including full length SHH:GFP, N-SHH:GFP and N-SHH when co-expressed with KIF17 (Figure 4I, Supplemental Figure 4G-H).

To investigate KIF17-mediated regulation of intracellular SHH levels *in vivo*, we employed an antibody directed toward the C-terminus of SHH [SHH antibody specificity was validated in P10 cerebella of *Shh* conditional mutant animals *Shh*^*CreER/lacZ*^ mice (Supplemental Figure 4I-Q)]. Intracellular SHH is detected in the Golgi/ER (horizontal arrowheads) and within the cell bodies of Purkinje cells of *Kif17*^*+/-*^ and *Kif17*^*-/-*^ littermates (vertical arrowheads, Figure 4J-M). However, SHH levels are significantly reduced in the posterior lobes of *Kif17*^*-/-*^ P10 cerebella (Figure 4N). Notably, SHH levels are not significantly altered in anterior lobes of *Kif17*^*-/-*^ mice (Supplemental Figure 4R). Together, these gain-and loss-of-function data suggest that KIF17 acts in Purkinje cells to stabilize intracellular SHH protein and promote SHH release. This is supported by the downregulation of HH target genes across the multiple HH-responsive cell types (CGNPs, BG and CGNs) following Purkinje cell-specific *Kif17* deletion. Reduction of SHH protein ultimately results in decreased CGNP proliferation and cerebellar hypoplasia in *Kif17* deletion mice (Figure 4O).

### *Kif17* deletion promotes CGNP proliferation *in vitro*

To investigate a role for KIF17 in CGNPs, we isolated and cultured wildtype and *Kif17*^*-/-*^ CGNPs *in vitro* (*30*). HH-dependent proliferation was measured in response to treatment with either Smoothened Agonist (SAG) or N-SHH conditioned media (N-SHH CM). Surprisingly, *Kif17*^*-/-*^ CGNPs display increased baseline proliferation compared to *Kif17*^*+/-*^ and *Kif17*^*+/+*^ CGNPs (Figure 5A-F). Treatment with either SAG or N-SHH CM resulted in increased CGNP proliferation, measured by EdU/BrdU incorporation (Figure 5E, Supplemental Figure 5A) or luminescence-based quantitation of ATP levels (Figure 5F). Additionally, we cultured CGNPs from *Kif17*^*fl/fl*^ and *Shh*^*Cre*^*;Kif17*^*fl/fl*^ littermates and evaluated their proliferation *in vitro* (Supplemental Figure 5B-G). We observed no significant differences of CGNP proliferation in *Kif17*^*fl/fl*^ and *Shh*^*Cre*^*;Kif17*^*fl/fl*^ cultures, confirming increased proliferation in *Kif17*^*-/-*^ CGNPs is a cell-autonomous phenotype. Notably, these results are distinct from those observed in CGNPs lacking *Boc*, which encodes for an essential HH co-receptor (*5*). Direct comparison of *Kif17*^*-/-*^ CGNP and *Boc*^*-/-*^ CGNP proliferation confirmed that *Kif17* deletion results in increased baseline and HH-stimulated CGNP proliferation (Supplemental Figure 5H).

**Figure 5.**
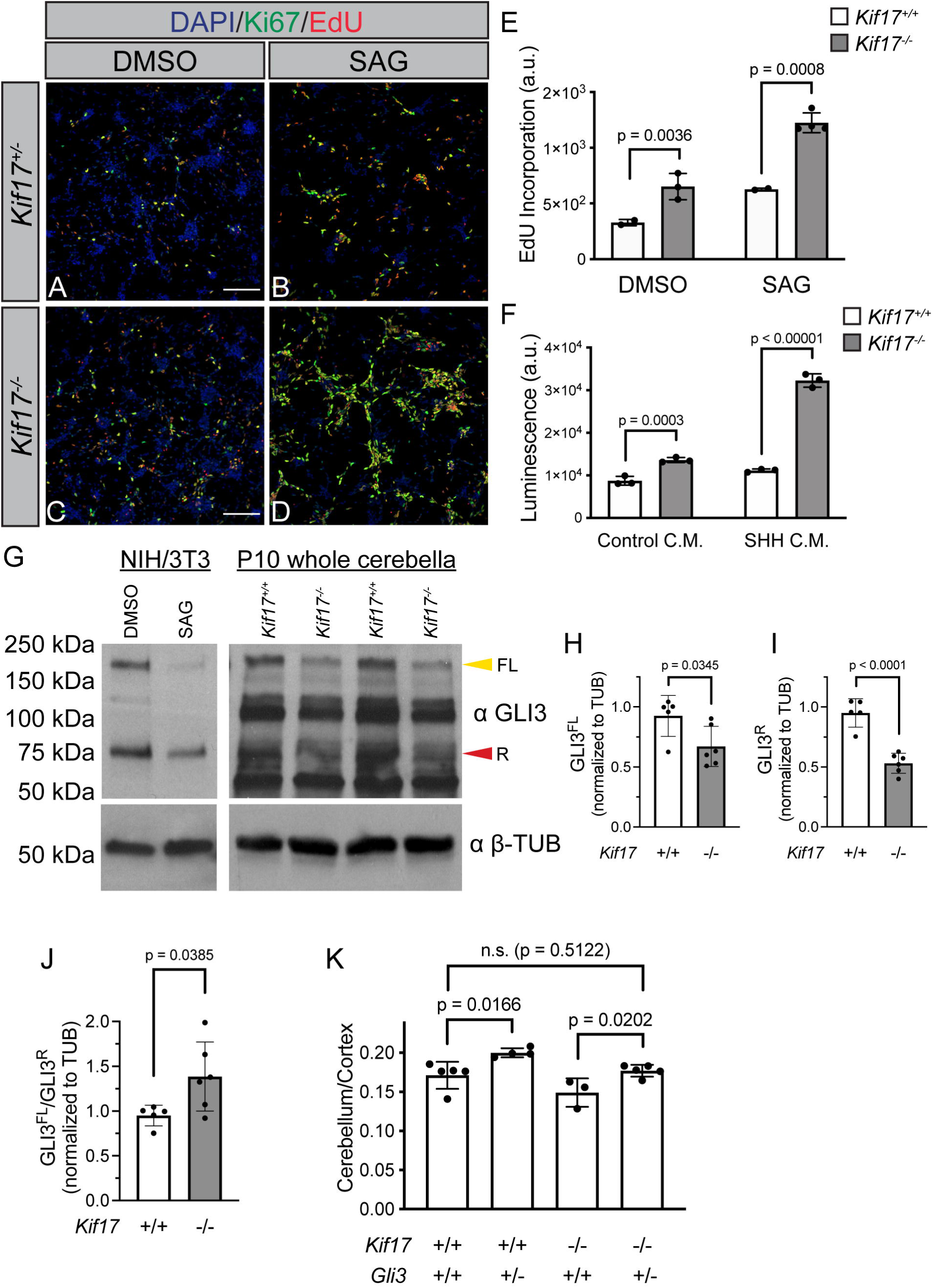
*Kif17* deletion promotes CGNP proliferation *in vitro*. Immunofluorescent analysis of proliferation in P8 CGNP cultures from *Kif17*^*+/-*^ (**A-B**) and *Kif17*^*-/-*^ (**C-D**) littermates. Antibody detection of Ki67 (green), fluorescent azide detection of EdU (red), and DAPI staining of nuclei (blue). Cultures were treated with DMSO as a vehicle control (**A, C**) or Smoothened agonist (SAG, **B, D**). Scale bars (**A, C**), 100 μm. Quantitation of EdU incorporation (**E**) in *Kif17*^*+/+*^ and *Kif17*^*-/-*^ CGNP cultures. Quantitation of ATP levels by luminescence values (**F**) in *Kif17*^*+/+*^ and *Kif17*^*-/-*^ CGNP cultures treated with control conditioned media (C.M.) or SHH conditioned media (SHH C.M.). Western blot analysis of GLI3 (**G**) in NIH/3T3 fibroblasts and P10 cerebella from *Kif17*^*+/+*^ and *Kif17*^*-/-*^ mice. The molecular masses (in kDa) of protein standards are indicated at the left of each blot. Yellow arrowhead denotes full length GLI3 (FL, 190 kDa); red arrowhead denotes GLI3 repressor (R, 83 kDa). Antibody detection of β-tubulin (⍰ β-TUB) was used to confirm equal loading across lanes. Quantitation of GLI3 full length (GLI3^FL^, **H**) and GLI3 repressor (GLI3^R^, **I**) normalized to β-tubulin. Ratio of GLI3^FL^ to GLI3^R^ (**J**) normalized to β-tubulin in *Kif17*^*+/+*^ and *Kif17*^*-/-*^ cerebella. Quantitation of cerebellar weight normalized to cortical weight in *Gli3;Kif17* compound mutants (**K**) at postnatal day 10. Data are mean ± s.d. Each dot represents an independent CGNP culture (**E-F**) or individual mouse (**H-K**). *P*-values were determined by a two-tailed Student’s *t*-test.

Given the altered baseline CGNP proliferation, we examined the levels and processing of the HH pathway transcriptional repressor, GLI3 in *Kif17* mutant animals. Western blot analysis of GLI3 full length (GLI3^FL^) and repressor (GLI3^R^) in P10 cerebella revealed (Figure 5G) significant reductions in both GLI3^FL^ and GLI3^R^ in *Kif17*^*-/-*^ cerebella (Figure 5H-I). Further, the ratio of GLI3^FL^ to GLI3^R^ is significantly increased in *Kif17* mutant cerebella (Figure 5J). These data suggest that similar to other kinesin-2 mutants (*31*), KIF17 regulates GLI3 processing in CGNPs. To examine the consequences of altering *Gli3* dosage in *Kif17* mutant animals, we measured cerebellar size in P10 *Kif17*^*lacZ*^*;Gli3*^*Xt*^ compound mutant cerebella (Figure 5K). Notably, loss of one *Gli3* allele causes cerebellar hyperplasia in *Kif17*^*+/+*^ mice and rescues the cerebellar hypoplasia phenotype observed in *Kif17* mutants. Together, these data suggest that KIF17 negatively regulates HH signaling in a cell-autonomous fashion within CGNPs, potentially through regulation of GLI3 repressor.

### CGNP-specific *Kif17* deletion results in a cell-autonomous HH gain-of-function phenotype

To directly assess KIF17 function in CGNPs *in vivo*, we crossed *Kif17*^*flox*^ mice to *Atoh1Cre* animals (Figure 6A), which specifically drives recombination in CGNPs [(*32*); Supplemental Figure 6A-F]. We used RT-qPCR to confirm efficient *Kif17* deletion in *Atoh1Cre;Kif17*^*fl/fl*^ cerebella (Figure 6B). Next, we assessed cerebellar size in *Atoh1Cre;Kif17*^*fl/fl*^ animals, which is unchanged compared to control animals (Figure 6C, Supplemental Figure 6G). These data are in striking contrast to *Kif17* germline mutants and Purkinje cell-specific *Kif17* deletion (cf. Figure 1S and Figure 3C). While thickness of the PCL is not significantly changed in either posterior or anterior lobes of *Atoh1Cre;Kif17*^*fl/fl*^ cerebella (Figure 6D, Supplemental Figure 6H), EGL thickness is increased, specifically in posterior lobes of *Atoh1Cre;Kif17*^*fl/fl*^ cerebella (Figure 6E, Supplemental Figure 6I). Notably, increased EGL thickness appears to be due to increased CGNP proliferation (as assessed by the percentage of EdU^+^ cells) in both posterior (Figure 6M) and anterior (Supplemental Figure 6K) lobes of *Atoh1Cre;Kif17*^*fl/fl*^ cerebella. RT-qPCR analysis revealed increased HH target gene expression in *Atoh1Cre;Kif17*^*fl/fl*^ cerebella compared to control littermates (Figure 6N-O; Supplemental Figure 6L-N). *In situ* hybridization confirmed that the increase in HH target gene expression is restricted to CGNPs in the posterior lobe, while no changes were observed in HH-responsive Bergmann glia and CGNs (Figure 6T; Supplemental Figure 6O). Together, these data indicate that CGNP-specific *Kif17* deletion results in increased HH pathway activity and CGNP proliferation, leading to a thicker EGL within posterior lobes of the developing cerebellum.

**Figure 6.**
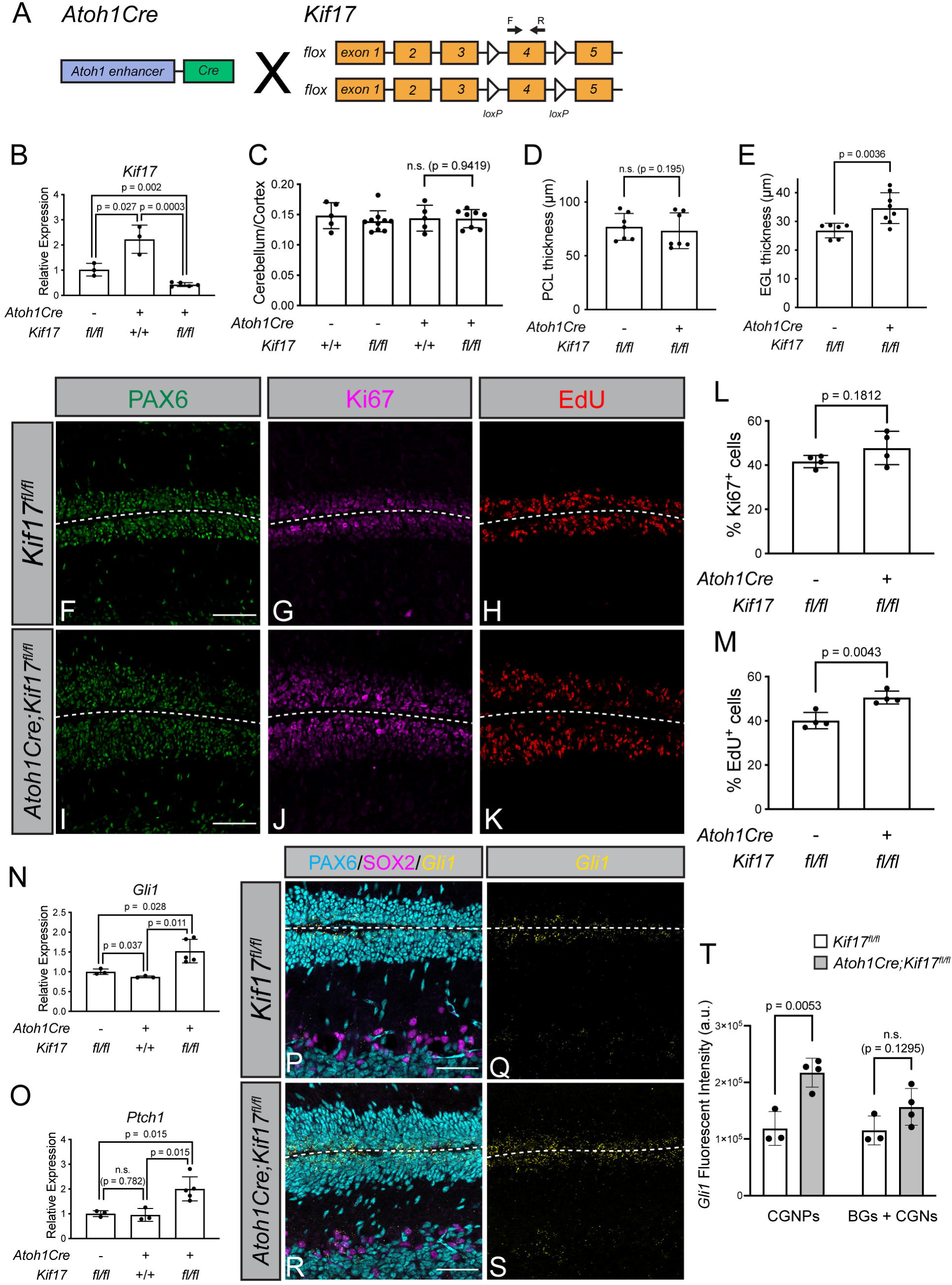
CGNP-specific *Kif17* deletion results in a cell-autonomous HH gain-of-function phenotype. (**A**) Schematic representing conditional *Kif17* deletion within CGNPs using *Atoh11Cre*. Arrows above exon 4 denote qPCR primers. Relative *Kif17* expression (**B**) measured by RT-qPCR in P10 cerebella from *Kif17*^*fl/fl*^, *Atoh1Cre;Kif17*^*+/+*^ and *Atoh1Cre;Kif17*^*fl/fl*^ mice. Cerebellar weights normalized to cortical weights (**C**) in P10 control and CGNP-specific *Kif17* deletion mice. Quantitation of PCL (**D**) and EGL (**E**) thickness in posterior lobes of *Kif17*^*fl/fl*^ and *Atoh1Cre;Kif17*^*fl/fl*^ P10 cerebella. Analysis of *in vivo* CGNP proliferation by immunofluorescence in *Kif17*^*fl/fl*^ (**F-H**) and *Atoh1Cre;Kif17*^*fl/fl*^ (**I-K**) P10 cerebella. Antibody detection of PAX6 (green; **F, I**) and Ki67 (magenta; **G, J**). Fluorescent azide detection of EdU (red; **H, K**). Scale bars (**F, I**), 50 μm. Percentage of Ki67^+^ (**L**) and EdU^+^(**M**) cells within the posterior cerebellar lobes of *Kif17*^*fl/fl*^ and *Atoh1Cre;Kif17*^*fl/fl*^ P10 mice. Relative *Gli1* (**N**) and *Ptch1* (**O**) expression measured by RT-qPCR in *Kif17*^*fl/fl*^, *Atoh1Cre;Kif17*^*+/+*^ and *Atoh1Cre;Kif17*^*fl/fl*^ P10 whole cerebella. Fluorescent *in situ Gli1* detection (yellow, **P-S**) with immunofluorescent detection of PAX6 to mark CGNPs and CGNs (cyan; **P, R**) and SOX2 to identify Bergmann glia (magenta; **P, R**) within posterior cerebellar lobes of P10 *Kif17*^*fl/fl*^ and *Atoh1Cre;Kif17*^*fl/fl*^ littermates. Scale bar (**P, R**), 50 μm. Quantitation of fluorescent intensity (integrated density) of *Gli1* puncta (**T**) within either CGNPs or Bergmann glia and cerebellar granule neurons (BGs + CGNs) in posterior cerebellar lobes of P10 *Kif17*^*fl/fl*^ and *Atoh1Cre;Kif17*^*fl/fl*^ mice. Data are mean ± s.d. Each dot represents an individual animal (**B-E, N-O**) or the average of 5 images per animal (**L-M, T**). *P*-values were determined by a two-tailed Student’s *t*-test. Dashed lines separate external granule layers.

### CGNP-specific *Kif17* deletion results in reduced GLI protein, increased CGNP proliferation, and elongated primary cilia *in vitro*

Given that other Kinesin-2 motors regulate GLI processing and trafficking, including in the cerebellum (*11, 12, 31*), we examined the consequences of CGNP-specific *Kif17* deletion on *Gli* expression and GLI protein levels. *Gli2* and *Gli3* expression are increased in *Atoh1Cre;Kif17*^*fl/fl*^ cerebella (Supplemental Figure 7A-B), similar to *Gli1*. However, western blot analysis (Figure 7A) revealed significantly reduced levels of GLI1 and GLI2 protein (Figure 7B-C). Similar to what was observed *Kif17*^*-/-*^ cerebella (cf Figure 5G-J), GLI3 full length and GLI3 repressor levels are also reduced (Figure 7D-E); further, the ratio of full length (GLI3^FL^) to repressor (GLI3^R^) is increased in *Kif17* mutant CGNPs (Figure 7F).

**Figure 7.**
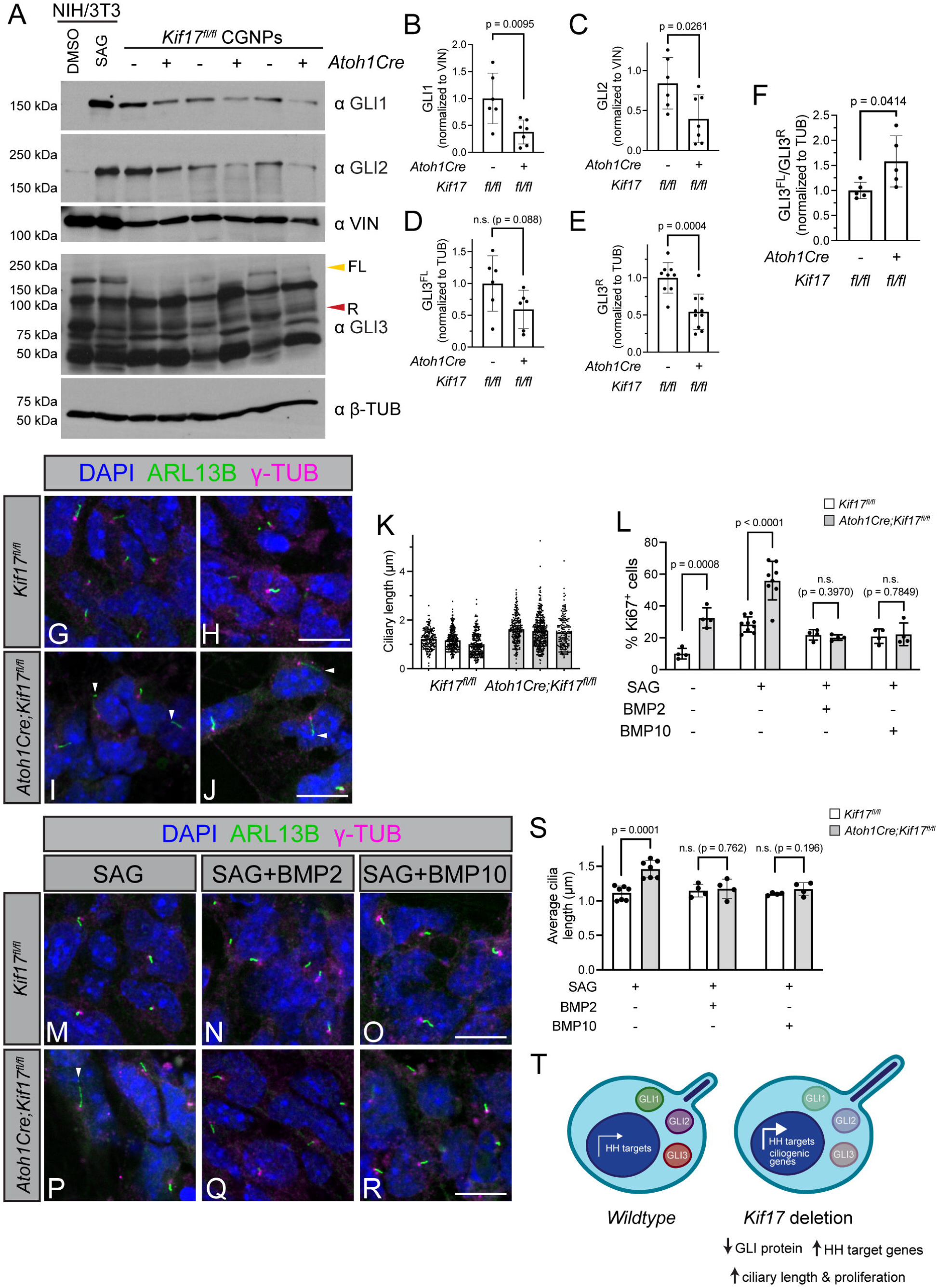
CGNP-specific *Kif17* deletion results in reduced GLI protein, increased CGNP proliferation, and elongated primary cilia *in vitro*. Western blot analysis (**A**) of GLI transcription factors (⍰ GLI1, ⍰ GLI2, ⍰ GLI3) in NIH/3T3 cells and purified CGNPs isolated from *Kif17*^*fl/fl*^ and *Atoh1Cre;Kif17*^*fl/fl*^ littermates. Antibody detection of Vinculin (⍰ VIN) confirmed equal loading across lanes for GLI1 and GLI2. Antibody detection of β-tubulin (⍰ β-TUB) confirmed equal loading across lanes for GLI3. Yellow arrowhead denote full length GLI3 (FL, 180 kDa) red arrowhead indicate GLI3 repressor (R, 83 kDa). The molecular masses (in kDa) of protein standards are indicated at the left of each blot. Quantitation of GLI1 (**B**) and GLI2 (**C**) normalized to Vinculin. Levels of GLI3^FL^ (**D**) and GLI3^R^ (**E**) normalized to β-Tubulin. Ratio of GLI3^FL^ to GLI3^R^ (**F**) in *Kif17*^*fl/fl*^ and *Atoh1Cre;Kif17*^*fl/fl*^ CGNPs, normalized to β-tubulin. Visualization of primary cilia (**G-J**) *in vitro* from *Kif17*^*fl/fl*^ (**G-H**) and *Atoh1Cre;Kif17*^*fl/fl*^ (**I-J**) CGNP cultures treated with SAG. Antibody detection of ARL13B (green) and γ-tubulin (γ-TUB, magenta) denote the axonemes and basal bodies of primary cilia, respectively; nuclei are identified with DAPI (blue). Scale bars (**H, J**), 10 μm. Ciliary length (**K**) quantitation of CGNPs from three representative cultures of *Kif17*^*fl/fl*^ and *Atoh1Cre;Kif17*^*fl/fl*^ littermates. Each dot represents an individual cilium. Percentages of Ki67^+^ (**L**) cells in CGNP cultures from P8 *Kif17*^*fl/fl*^ and *Atoh1Cre;Kif17*^*fl/fl*^ littermates. Cells were treated with vehicle (DMSO) or SMO Agonist (SAG), with or without BMP2 or BMP10. Immunofluorescent detection of primary cilia *in vitro* from *Kif17*^*fl/fl*^ (**M-O**) and *Atoh1Cre;Kif17*^*fl/fl*^ (**P-R**) CGNP cultures treated with SAG and BMP ligands. Antibody detection of ARL13B (green) and γ-Tubulin (γ-TUB, magenta) label the axonemes and basal bodies of primary cilia; nuclei are identified with DAPI (blue). Scale bar (**O, R**), 10 μm. Average CGNP ciliary length (**S**) from *Kif17*^*fl/fl*^ and *Atoh1Cre;Kif17*^*fl/fl*^ littermates, where each dot represents the average length of an individual culture. Data are mean ± s.d. *P*-values were determined by a two-tailed Student’s *t*-test. (**T**) Summary of CGNP-specific *Kif17* deletion on GLI proteins, CGNP proliferation and primary cilia length.

We also assessed potential physical interactions between KIF17 and GLI proteins, as previously demonstrated for other Kinesin-2 motors (*13*). Co-immunoprecipitation of epitope-tagged KIF17 (KIF17:HA) and GLI transcription factors (MYC:GLI1, MYC:GLI2, MYC:GLI3) suggested that KIF17 can indeed physically interact with all three GLI proteins (Supplemental Figure 7C).

We noted that *Atoh1* expression is increased in animals with CGNP-specific *Kif17* deletion (Supplemental Figure 7D); previous work demonstrated that ATOH1 promotes ciliogenesis and maintains CGNP responsiveness to HH (*33*). However, analysis of CGNP primary cilia length in *Atoh1Cre;Kif17*^*fl/fl*^ P10 cerebella revealed no significant change *in vivo* (Supplemental Figure 7E-F; p = 0.4534 for posterior lobes, p = 0.0886 for anterior lobes). In contrast, when we examined primary ciliary length in SAG-treated CGNPs *in vitro*, we found that CGNPs lacking *Kif17* display increased ciliary length (Figure 7G-L), with an average ciliary length of 1.46 μm (compared to 1.1 μm in control animals); notably, some primary cilia reached lengths of 5 μm (Figure 7K).

Since HH signaling also regulates cilia length (*34*) and ciliogenesis (*35*), we investigated whether increased ciliary length was a cause or a consequence of HH pathway activity. We antagonized HH signaling *in vitro* by adding BMP ligands, either BMP2, which has been previously shown to antagonize SHH-induced CGNP proliferation (*36*) or BMP10, which is significantly upregulated in *Kif17*^*-/-*^ cerebella (Supplemental Figure 7G). Notably, both BMP2 and BMP10 effectively attenuate HH-mediated CGNP proliferation in both *Kif17*^*fl/fl*^ and *Atoh1Cre;Kif17*^*fl/fl*^ cultures (Supplemental Figure 7H-P, Figure 7L). However, BMP2 and BMP10 treatment reduced ciliary length specifically in *Atoh1Cre;Kif17*^*fl/fl*^ CGNPs (Figure 7M-S, Supplemental Figure 7Q), resulting in average ciliary lengths of 1.18 μm (BMP2) and 1.17 μm (BMP10). Together, these data suggest that high levels of HH pathway activation in *Kif17* mutant CGNPs results in increased ciliary length, which can be attenuated by BMP signaling.

## Discussion

In this study we investigated a role for the kinesin-2 motor KIF17 in HH-dependent cerebellar development. Our work revealed that *Kif17* is expressed in both SHH-producing Purkinje cells and SHH-responsive CGNPs. Purkinje cell-specific *Kif17* deletion phenocopies germline *Kif17* deletion, resulting in reduced EGL thickness due to reduced HH target gene expression and decreased CGNP proliferation. Conversely, CGNP-specific *Kif17* deletion increased EGL thickness due to increased HH target gene expression and increased CGNP proliferation (Figure 8). This work identifies dual and opposing roles for KIF17 in HH-dependent cerebellar development– first, as a positive regulator of HH signaling through regulation of SHH protein levels within Purkinje cells, and second, as a negative regulator of HH signaling through regulation of GLI transcription factors in CGNPs.

**Figure 8.**
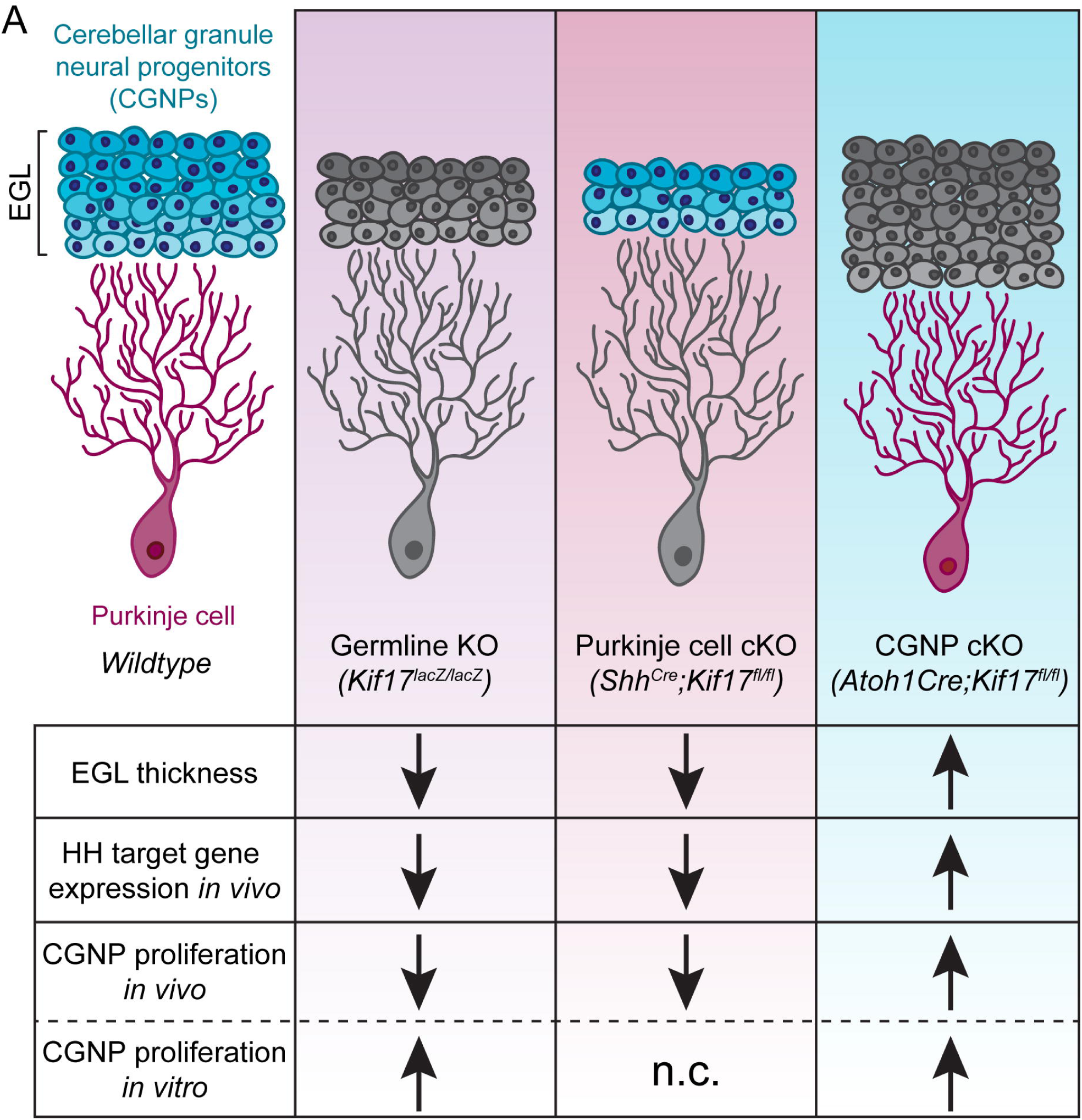
KIF17 has dual and opposing roles in HH signaling in the developing cerebella. (**A**) Schematic demonstrating the consequences of KIF17 deletion on cerebellar development. Germline or Purkinje cell-specific *Kif17* deletion (magenta) results in a HH loss-of-function phenotype *in vivo*, while CGNP-specific *Kif17* deletion yields a HH gain-of-function phenotype (cyan). Notably, germline *Kif17* deletion results in increased CGNP proliferation *in vitro*, similar to CGNP-specific *Kif17* deletion both *in vivo* and *in vitro*.

### KIF17 function in SHH-producing Purkinje cells

Here we demonstrated that KIF17 is required in Purkinje cells to mediate proper HH-dependent cerebellar development and that KIF17 regulates SHH protein levels within Purkinje cells. Specifically, we visualized intracellular SHH utilizing a C-terminal antibody, which revealed reduced SHH protein in *Kif17* mutant cerebella, both within the presumed endoplasmic reticulum/Golgi apparatus and more broadly within Purkinje cell bodies. Notably, SHH is translated as a 45 kDa precursor protein, which undergoes auto-catalytic cleavage into a 19 kDa N-terminal fragment and 25 kDa C-terminal fragment (*37-39*). The N-terminal fragment is dually-lipidated with cholesterol at the C-terminus and palmitate at the N-terminus to produce active ligand (reviewed in (*40*)). While the 25 kDa C-terminal SHH fragment does not transduce HH signaling, the C-terminal HH fragment does target N-HH to axons and growth cones in the developing retina of *Drosophila melanogaster* (*41*). One model for KIF17 action in Purkinje cells is the transport of SHH-containing vesicles along microtubules to distinct locations within these cells. This model has precedence with a previously described role for KIF17 in the vesicular trafficking of NR2B in the hippocampus (*17, 25*). Further, this is consistent with the reduced levels of SHH protein in *Kif17* mutants, as NR2B levels are also reduced when its vesicular trafficking is disrupted in *Kif17* mutants. This model is also consistent with the results from KIF17 gain-of-function experiments demonstrating increased intracellular SHH protein accumulation (this study). However, we cannot rule out similar trafficking-related effects of KIF17 on other HH pathway components, such as SCUBE2 and DISP, both of which regulate SHH protein release from cell surfaces. We also cannot distinguish between KIF17-mediated effects on SHH trafficking versus potential impacts on SHH protein stability. Distinguishing between these possibilities would require robust methods to culture Purkinje cells *ex vivo*, which are currently lacking.

### KIF17 regulation of GLIs in CGNPs

In addition to a non-cell autonomous role for KIF17 in Purkinje cells, we also established a cell autonomous role for KIF17 in CGNPs, where *Kif17* deletion results in a HH gain-of-function phenotype– increased CGNP proliferation and upregulation of several HH target genes. CGNP-specific *Kif17* deletion results in reduced protein levels of all three HH transcriptional effectors, GLI1, GLI2, and GLI3. Previous work established GLI1 and GLI2 as transcriptional activators in the developing cerebellum where *Gli2* deletion results in a HH loss-of-function phenotype (*4, 6*). Given these roles for GLI1 and GLI2, we were surprised to find that CGNP-specific *Kif17* deletion results in a HH gain-of-function phenotype. However, the concomitant loss of GLI3 repressor in *Kif17* mutant CGNPs suggests that GLI repressor function is a significant mediator of CGNP proliferation. Notably, reduction of GLI activator and repressor protein is consistent with previous work where cerebellar-specific Suppressor of fused (*Sufu*) deletion also results in increased CGNP proliferation (*42*). GLI3 also acts during early embryonic cerebellar development in mesencephalon and rhombomere 1 patterning through the regulation of *Fgf8* expression (*43*). Here we also show that loss of one *Gli3* allele is sufficient to drive cerebellar hyperplasia, likely due to increased HH signaling. Together, these data suggest that KIF17 in CGNPs promotes GLI3 repressor formation to restrict proliferation in the postnatal cerebellum, consistent with previous work demonstrating central roles for other kinesin-2 motors in GLI processing (*11, 31, 44*).

GLIs require primary cilia for proper processing and transcriptional activity (reviewed in (*8*)). Further, studies have established ciliary tip localization of KIF17 (*45*), similar to GLI transcription factor localization during HH activation (*46, 47*). One model for KIF17 regulation of GLI protein levels in CGNPs is through ciliary trafficking or localization. Notably, this is consistent with recent work demonstrating that GLI interactions with KIF7 promote ciliary localization (*48*). Unfortunately, the lack of suitable KIF17 antibodies precludes rigorous testing of this hypothesis. Other possible roles for KIF17 in CGNPs include the regulation of GLI trafficking and stability as well as interactions with other ciliary proteins that regulate GLI processing, such as SUFU, KIF7 or PKA (*49-51*).

### Kinesin motors and HH signaling

While previous studies have explored the requirement for kinesin and dynein motors in HH-responding cells (reviewed in (*8*)), the current study highlights a novel role for kinesin motors in HH-producing cells. An outstanding question is whether KIF17 functions in HH-producing cells in other tissues. Notably, the subgranular zone of the hippocampus and subventricular zone rely on proper HH signaling for neurogenesis (*52-55*). While KIF17 has a well-defined role in NR2B trafficking in the hippocampus (*17, 25*), the potential contribution of KIF17 to HH signaling in the hippocampus has not yet been examined.

In addition to its neural-specific contributions, KIF17 has several described functions in the testes, although loss-of-function studies have yet to be performed (*56-60*). Desert Hedgehog (DHH) is expressed in Sertoli cells, and *Dhh* deletion results in a loss of HH-responsive Leydig cells (*61*). While we did not observe infertility in *Kif17* mutant mice, it will be of interest to investigate the consequences of *Kif17* deletion on HH-dependent spermatogenesis. Future studies investigating the contribution of other kinesin-2 motors, particularly KIF3A/KIF3B, in HH-producing cells (e.g., in the notochord or zone of polarizing activity) will be of high interest. Finally, this work raises the question of potential contributions from KIF3C, another accessory kinesin-2 motor, to HH signal transduction.

## Materials and Methods

### Animal models

*Kif17*^*lacZ*^ germline mutant mice have been previously described (*22*). These mice were maintained on two different congenic C57BL/6J and 129S4/SvJaeJ backgrounds after backcrossing for at least 10 generations. *Kif17*^*fl*^ animals carrying *Kif17* conditional alleles were generated from the initial knock-in allele from EUCOMM through crossing *Kif17*^*tm1A*^ animals to ubiquitous Flippase mice obtained from The Jackson Laboratory [strain 011065, (*62*)] to generate *Kif17*^*tm1C*^*/Kif17*^*flox*^ mice. These mice were maintained on a congenic C57BL/6J background. *Atoh1Cre* animals were obtained from The Jackson Laboratory [strain 011104, (*32*)] and maintained on a C57BL/6J background. Mice carrying the *Shh*^*Cre*^ allele [strain 005622] were provided by Dr. Deb Gumucio and previously described (*26*). These mice were backcrossed for at least 10 generations to C57BL/6J animals to create a congenic line. All animal procedures were reviews and approved by the Institutional Animal Care and Use Committee (IACUC) at the University of Michigan, USA.

### Wholemount X-gal staining

Postnatal cerebella were dissected in 1X PBS (pH 7.4) and cut in half with a razor before fixation (1% formaldehyde, 0.2% glutaraldehyde, 2 mM MgCl_2_, 5 mM EGTA, 0.02% NP-40) on ice for 20 min. After fixation, the cerebella were washed 3 × 5 min with 1X PBS (pH 7.4) on a rocking platform. Beta-Galactosidase activity was detected with X-gal staining solution [5 mM K_3_Fe(CN)_6_, 5 mM K_4_Fe(CN)_6_, 2 mM MgCl_2_, 0.01% Na deoxycholate, 0.02% NP-40, 1 mg/ml X-gal]. The signal was developed for 24 h at 37°C, changing the staining solution after 12 h. After staining, cerebella were washed 3 × 5 min with 1X PBS (pH 7.4) and post-fixed in 4% paraformaldehyde for 30 min at room temperature on a rocking platform, followed by 3 × 5 min washes in 1X PBS (pH 7.4). Cerebella were photographed using a Nikon SMZ1500 microscope and stored in 1X PBS (pH 7.4).

### Section Immunofluorescence

Section immunofluorescence was performed as described in (*63*). Briefly, cerebella were dissected in 1X PBS (pH 7.4) and cut in half using a razor. For all experiments except for beta-galactosidase and SHH visualization, cerebella were fixed with 4% paraformaldehyde (Electron Microscopy Sciences) for 1 h on ice. For beta-galactosidase immunofluorescence, cerebella were fixed (1% formaldehyde, 0.2% glutaraldehyde, 2 mM MgCl_2_, 5 mM EGTA, 0.02% NP-40) on ice for 20 min. For SHH visualization, cerebella were fixed in Sainte Marie’s solution (95% ethanol, 1% acetic acid) at 4°C on a rocking platform for 24 h. Following fixation, cerebella were washed 3 × 5 min with 1X PBS (pH 7.4) on a rocking platform and cryoprotected overnight in 1X PBS + 30% sucrose on a rocking platform. Then, cerebella were washed 3 × 1 h in 50% OCT (Fisher Scientific, 23-730-571) before embedding in 100% OCT. Sections were collected on a Leica CM1950 cryostat at 12 μm thickness for all experiments, except for SHH visualization, which were sectioned at 9 μm thickness. Slides were then washed 3 × 5 min with 1X PBS (pH 7.4). For mouse primary antibodies, citric acid antigen retrieval (10 mM citric acid + 0.5% Tween-20, pH 6.0) at 92°C for 10 min was performed prior to primary antibody incubation. Primary antibodies were diluted in blocking buffer (3% bovine serum albumin, 1% heat-inactivated sheep serum, 0.1% Triton X-100) and incubated overnight at 4°C in a humidified chamber. After primary antibody incubation, slides were washed 3 × 10 min with 1X PBST^X^ (1X PBS + 0.1% Triton X-100, pH 7.4). Secondary antibodies were diluted in blocking buffer and incubated for 1 h at room temperature, followed by 3 × 5 min 1X PBST^X^ washes. Nuclei were labeled using DAPI (0.5 μg/mL in blocking buffer) for 10 min and washed twice with 1X PBS. Coverslips were mounted using Immu-mount aqueous mounting medium (Thermo Fisher Scientific, 9990412). Images were taken on a Leica SP5X upright confocal (2 photon). A list of all the primary and secondary antibodies and their working concentrations is provided in Table S1.

### Fluorescent *in situ* hybridization

Cerebella were dissected in 1X PBS (pH 7.4) and cut in half using a razor. Cerebella were fixed with 10% neutral buffered formalin (Fisher, 245-685) on a rocking platform at room temperature for 24 h. Following fixation, cerebella were washed 3 × 5 min with 1X PBST^X^ on a rocking platform and cryoprotected overnight in 1X PBS + 30% sucrose on a rocking platform. Cerebella were then washed 3 × 1 h with 50% OCT compound before embedding in 100% OCT. Sections were collected on a Leica CM1950 cryostat at 12 μm thickness. Slides were processed using RNAscope Multiplex Fluorescent Detection kit (ACD, 323110) using a protocol adapted from (Holloway, 2021). Prior to probe hybridization, samples underwent antigen retrieval for 15 minutes and treated with Protease Plus (ACD, 322381) for 5 minutes. Probes used in this paper were *Mm*-*Gli1* (ACD, 311001) and *E*.*coli-lacZ* (ACD, 313451). After probe detection, slides were subsequently stained using the above-described section immunofluorescence protocol.

### RT-qPCR

Cerebella were dissected in 1X PBS, and RNA was isolated using a PureLink RNA Mini Kit (ThermoFisher Scientific, 12183025). Following isolation, 2 μg of RNA were used to generate cDNA libraries using a High-Capacity cDNA reverse transcription kit (Applied Biosystems, 4368814). RT-qPCR was performed using PowerUP SYBR Green Master Mix (Applied Biosystems, A25742) in a QuantStudio 3 Real-Time PCR System (Applied Biosystems). Primers used in this paper can be found in Table S2. Gene expression was normalized to *Gapdh*, and relative expression analyses were performed using the 2(^-ddCT^) method. For RT-qPCR analysis, biological replicates were analyzed in triplicate.

### Weight analyses

For weight measurements, the date litters were born were noted as postnatal day 0 and were dissected on postnatal day 10. Pups were first weighed and then placed on ice briefly before decapitation. The cortices and cerebella were dissected in 1X PBS (pH 7.4). To weigh cortices and cerebella, a specimen jar was first filled with PBS on an analytical scale. The tissue was transferred with forceps to the specimen jar, and its weight was recorded. Genotyping samples were taken after dissection, allowing the weights to be recorded without prior knowledge of the genotype.

### EGL and PCL thickness quantitation

To measure the thickness of the external granule layer (EGL) and Purkinje cell layer (PCL), ImageJ software was utilized. Images were first blinded before measuring. For EGL thickness, the measurement was taken to include the top of the nuclei at the uppermost position in the EGL to the bottom of the nuclei at the bottommost position in the EGL. For PCL thickness, the measurement was taken just below the bottommost nuclei in the EGL to the center of Purkinje cell nuclei within the molecular layer. For each image, at least 10 measurements were obtained. For each animal, at least three images were acquired in the posterior lobes and an additional three images in the anterior lobes.

### EdU incorporation assay *(in vivo)*

On postnatal day 9, pups were intraperitoneally injected with 100mg/kg of EdU (Invitrogen, A10044), dissolved in 1X PBS (pH 7.4). 24 h later, cerebella were dissected and processed for section immunofluorescence as described above. Prior to primary antibody incubation, EdU incorporation was visualized with an azide staining solution [100 mM Tris HCl (pH 8.3), 0.5 mM CuSO_4_, 50 mM ascorbic acid, 50 μM Alexa Fluor 555 Azide, Triethylammonium Salt (Thermo Fisher Scientific, A20012)] for 30 min at room temperature. Sections were then washed 3 × 10 min in PBST^X^ (1x PBS + 0.1% Triton X-100, pH 7.4), followed by immunofluorescence staining as described above.

### Section digoxigenin *in situ* hybridization

Section digoxigenin *in situ* hybridization was performed as previously described (Allen at al., 2011; Wilkinson, 1992). First, cerebella were dissected in 1X PBS (pH 7.4) and fixed for 24 h with 4% paraformaldehyde at 4°C on rocking platform. After fixation, cerebella were washed 3 × 5 min with 1X PBST^W^ (1X PBS + 0.1% Tween-20, pH 7.4) and cryoprotected with 1X PBS + 30% sucrose overnight on a rocking platform. The next day, cerebella were subjected to 3 × 1 h washes with 50% OCT before embedding in 100% OCT. Cerebella were sectioned on Leica CM1950 cryostat at 20 μm thick sections. Probe hybridization was performed with the indicated digoxigenin probes at a concentration of 1 ng/μl overnight at 70°C. The sections were incubated in AP-conjugated anti-DIG antibody (Table S1). AP-anti-DIG was visualized with BM Purple (Roche, 11442074001), and signal was developed for 4 h at 37°C. After the signal was developed, development was stopped with 3 × 5 min washes with 1X PBS (pH 4.5). Sections were post-fixed in 4% PFA + 0.2% glutaraldehyde for 30 min, then washed 3 × 5 min in 1X PBS (pH 7.4). Sections were dried with 70% ethanol wash before drying at 60°C for 10 min. Coverslips were mounted using Glycergel (DAKO, C056330-2) preheated to 60°C. Images were taken on a Nikon SMZ1500 microscope.

### Western blot analysis

#### For cerebellar lysates

Cerebella were dissected in 1X PBS (pH 7.4) and lysed in radioimmunoprecipitation assay buffer [50 mM Tris-HCl (pH 7.2), 150 mM NaCl, 0.1% Triton X-100, 1% sodium deoxycholate, 5 mM EDTA] containing protease inhibitor (Roche, 11836153001) and 1 mM phenylmethylsulfonyl fluoride (PMSF; Sigma, 10837091001). Extracts were cleared by centrifugation at 21130 rcf for 10 min at 4°C. Total protein concentration was determined with the Pierce BCA protein assay kit (Thermo Fisher Scientific), utilizing 50 μg of cerebellar lysate for each sample. Lysates were mixed with 6X Laemmli buffer and denatured at 95°C for 10 min. Protein was separated by SDS-PAGE (5% separating gel for GLI1 and GLI2, 6.25% for GLI3 and 12% for SHH) and transferred onto Immuno-Blot PVDF membranes (Bio-Rad) at 100 v for 100 min on ice. For most blots, primary antibodies were diluted in blocking buffer [30 g/L bovine serum albumin with 0.2% NaN_3_ in 1X TBST (Tris-buffered saline, 0.5% Tween-20, pH 7.4)]. Blots were incubated with primary antibodies overnight at 4°C on a rocking platform. For detecting SHH, the primary antibody was diluted in 1X TBST and blots were incubated with primary antibody for 1 h at room temperature on a rocking platform. All primary antibodies and concentrations used can be found in Table S1. After incubation with primary antibody, blots were washed 3 × 10 min in 1X TBST. Peroxidase-conjugated secondary antibodies (Table S1) were diluted in blocking buffer, and blots were incubated with secondary antibodies for 1 h at room temperature on a rocking platform. After secondary incubation, blots were washed 4 × 10 min in 1X TBST, following incubation with Amersham ECL Prime Western Blotting Detecting Reagent (GE Healthcare, RPN2232) for 2 min and then exposed to HyBlot CL autoradiography film (Fisher Scientific, NC9556985) and developed using a Konica Minolta SRX-101A medical film processor. Relative levels were obtained by taking the integrated density value of each band, subtracting the background of the lane, and normalizing to the integrated density of housekeeping protein (CALB-1, VINCULIN, β-TUB) minus the background of the lane.

#### For Cos7 overexpression lysates

COS-7 cells were transiently transfected with the relevant DNA constructs using Lipofectamine 2000 (Invitrogen, catalog number 11668). The media was collected, and cells were lysed 48 h after transfection in HEPES lysis buffer (25 mM HEPES pH 7.4, 115 mM KOAc, 5 mM NaOAc, 5 mM MgCl2, 0.5 mM EGTA and 1% Triton X-100) containing protease inhibitor (Roche, catalog number 11836153001) and 1 mM PMSF (Sigma, 10837091001). Culture media and extracts were cleared by centrifugation at 15,000 rpm for 10 min at 4°C. Total protein concentration was determined for cell lysates with the Pierce BCA protein assay kit (Thermo Fisher Scientific), utilizing 50 μg of cell lysate for each sample. Collected culture media was diluted 1:5 in HEPES lysis buffer before mixing with 6X Laemmli buffer and denatured at 95°C for 10 min. Protein was separated by SDS-PAGE using 12% gels and transferred onto Immuno-Blot PVDF membranes (Bio-Rad). Membranes with cell lysates were treated identical to cerebellar lysates, as described above.

### Tamoxifen induction

To conditionally delete *Shh* in *Shh*^*CreER/lacZ*^ mice, neonatal pups were injected intraperitoneally with 50 mg/kg of tamoxifen (Sigma, T5648-1G) dissolved in corn oil once daily on postnatal days 7, 8 and 9. On postnatal day 10, cerebella were collected and processed for section immunofluorescence described above.

### Cerebellar granule neuronal progenitor cultures

The protocol was adapted from (*30*). Postnatal day 8 animals were anesthetized on ice briefly before decapitation. Cerebella were dissected in 1X PBS (pH 7.4) and placed in Hibernate-A media (BrainBits, HA). Tissue was then washed once with 1X PBS (pH 7.4). Cerebella were incubated in digestion media [0.25% Trypsin-EDTA (Gibco, ILT25200056) + 1 mg/mL DNAse I (Roche, 10104159001)] for 5 min at 37°C followed by trituration with a P1000 pipette, and subsequent incubation for 15 min at 37°C, shaking the dish every 5 minutes. After digestion, the pieces of tissue were further broken up with a P1000 pipette and transferred to a conical containing isolation media [DMEM (Gibco, 11965-092) + 10% calf bovine serum (ATCC 50-189-025NP) + 1x Penicillin-Streptomycin-Glutamine (Gibco, 10378016)]. The digested tissue was spun down 800 rcf for 8 min to pellet the cells. Digestion media was removed, and the pellet was washed with twice more isolation media. The pellet was fully resuspended in isolation media and passed through a 70 μm cell strainer. Single cell suspensions were spun down and resuspended in 1 mL of isolation media, which was then added to the top of a 30%/60% Percoll gradient (Sigma/Cytiva, P1644) before spinning at 800 rcf for 20 min. Initially, 100% Percoll was diluted with 10X PBS to make 90% Percoll. For 60% Percoll, 90% Percoll was diluted in L15 complete media [Leibovitz’s L-15 Medium without phenol red (Gibco, 21083027) + 10% calf bovine serum (ATCC 50-189-025NP) + 1x Penicillin-Streptomycin-Glutamine (Gibco, 10378016)]. For 30% Percoll, 90% Percoll was diluted in isolation media. CGNPs were isolated from the 30%/60% Percoll interphase and washed with isolation media. Finally, CGNPs were resuspended in neuronal media [Neurobasal media (Gibco, 21103049) + 1% calf bovine serum (ATCC 50-189-025NP) + 1x Penicillin-Streptomycin-Glutamine (Gibco, 10378016) + 1x B27 supplement (Gibco, 17504044)] and counted using a hemocytometer and plated at appropriate densities onto chambers or wells that were incubated with laminin (Sigma, L2020). CGNPs were cultured at 37°C, 5% CO2, 95% humidity in neuronal media. For activation of Hedgehog signaling, either SHH C.M. collected from COS-7 cells was added to the media (1:10) or 500 nM of SAG (Enzo Life Sciences, ALX-270-426-M001) dissolved in DMSO was added to the media. To antagonize HH signaling, BMP2 (Peprotech, 120-02) was used at 100 ng/mL, and BMP10 (Peprotech, 120-40) was used at 10 ng/mL. Half media changes were done every 24 hours for the duration of the cultures. 24 h before fixation, 10 μM EdU (Invitrogen, A10044), dissolved in DMSO, was administered to the culture.

### Genotyping with beta-galactosidase fluorescence

For co-culturing *Kif17*^*+/-*^ and *Kif17*^*-/-*^ CGNPs, we utilized BetaFluor β-gal assay kit (Promega 70979-3) to distinguish between *Kif17*^*+/-*^ and *Kif17*^*-/-*^ littermates at postnatal day 8. Briefly, while dissected cerebella were on ice in Hibernate-A media, half of the cortex placed in TrypLE express (Invitrogen, ILT12604013) for 15 minutes at 37°C before lysing with reporter lysis buffer (Promega, E397A). Samples were spun at 15,000 rpm for 10 min at 4°C, and supernatant removed to a fresh tube. Lysates were then plated in triplicate in clear bottom 96-well plate and incubated with assay mixture for 30 min at 37°C before reading fluorescence. Genotyping samples taken at dissection later confirmed beta-galactosidase assay results.

### Cerebellar granule neuronal progenitor culture immunofluorescence

Culture media was removed gently before coverslips were fixed in 4% paraformaldehyde for 30 min at room temperature. Coverslips were washed 3 × 5 min with 1X PBST^X^, then were stained with EdU staining solution [100 mM Tris HCl (pH 8.3), 0.5 mM CuSO_4_, 50 mM ascorbic acid, 50 µM Alexa Fluor 555 Azide, Triethylammonium Salt (Thermo Fisher Scientific, A20012)] for 30 min at room temperature. Coverslips were washed 3 × 5 min with 1X PBST^X^ and then blocked with blocking buffer (3% bovine serum albumin, 1% heat-inactivated sheep serum, 0.1% Triton X-100) for either 1 h at room temperature or 4°C overnight. Primary antibodies were diluted in blocking buffer. Coverslips were removed from the plate and were placed onto the diluted primary antibodies on top of parafilm for 1 h at room temperature. Coverslips were placed back in the well and were washed 3 × 5 min with 1X PBST^X^. Secondary antibodies were diluted in blocking buffer and were added to a fresh piece of parafilm. Coverslips were placed onto the parafilm and incubated with the secondaries for 1 h at room temperature. After secondary incubation, nuclei were labeled using DAPI (0.5 ng/mL in block buffer) for 10 minutes. Coverslips were then washed 3 × 5 min with PBST^X^. Before mounting onto a slide with Immu-mount aqueous mounting medium (Thermo Fisher Scientific, 9990412), coverslips were briefly dipped in water. Images were taken on a Leica SP5X upright confocal (2 photon).

### CGNP microplate assays

To quantify EdU incorporation *in vitro*, a Click-iT EdU proliferation assay (Thermo Fisher Scientific, C10499) was used in CGNPs *in vitro*. 24 h after plating, EdU was added to the culture (10μM, dissolved in DMSO). 48 h after plating, the assay was completed according to the manufacturer’s protocol. To measure BrdU incorporation, a colorimetric BrdU Cell Proliferation ELISA Kit (Abcam, ab126556) was utilized. 48 h after plating (2 d *in vitro*), BrdU was administered to the culture. 48 h after BrdU addition (4 d *in vitro*), the assay was completed according to the manufacturer’s protocol. To quantify the number of viable CGNPs *in vitro*, a CellTiter-Glo® Luminescent Cell Viability Assay (Promega, G7570) was used on cultures grown for 4 d *in vitro*. The assay was performed according to the manufacturer’s protocol.

### Immunoprecipitation of tagged proteins

COS-7 cells were transiently transfected with the relevant DNA constructs using Lipofectamine 2000 (Invitrogen, 11668). Cell lysates (1 mg) were pre-cleared with Protein-G–agarose beads (Roche, catalog number 11719416001) for 1 h at 4°C. MYC-or HA-tagged proteins were immunoprecipitated from pre-cleared lysates using either anti-MYC or anti-HA antibodies for 2 hours at 4°C. Following immunoprecipitation, the lysates were incubated with Protein-G–agarose beads for 1 h at 4°C. The Protein-G– agarose beads were subjected to 5 × 8 min washes in HEPES lysis buffer and resuspended in 30 μl of 1X PBS and 6X Laemmli buffer. The samples were boiled for 10 min and proteins were separated using SDS-PAGE and analyzed by western blotting. Visualization and quantitation were identical to the above-described western blot analysis.

### Image quantitation

To quantify intensity of SHH immunofluorescent signal, ImageJ software was used to measure the fluorescence integrated density of individual Purkinje cell bodies, subtracting the background measured from the internal granule layer. Per mouse, at least 5 images from the posterior lobes were measured, and an additional 5 images of the anterior lobes. To quantify fluorescent *Gli1* fluorescence, ImageJ software was used to measure the integrated density fluorescent signal contained to either the external granule layer (EGL, CGNPs) or lower molecular layer to inner granule layer (IGL, Bergmann glia and CGNs). At least six images were analyzed per mouse; three images for each posterior and anterior lobes. For all image analyses, images were blinded.

### Quantitation and statistical analysis

All the data are mean ± s.d. All statistical analyses were performed using GraphPad Prism (www.graphpad.com). Statistical significance was determined by using a two-tailed Student’s *t*-test. For all the experimental analyses, a minimum of three mice of each genotype were analyzed, each *n* represents a mouse. For *in vitro* experiments, a minimum of three biological replicates were analyzed, each *n* represents a biological replicate. All the statistical details (statistical test used, adjusted *P*-value, statistical significance and exact value of each *n*) for each experiment are specified in the figure legends.

## Acknowledgments

We thank past and present Allen lab members for their valuable feedback and suggestions. We also thank Joe Besharse (Medical College of Wisconsin) for providing the *Kif17* mutant mice. We thank Suzie Scales (Genentech) and Martin Engelke (Illinois State University) for their insightful comments. We thank Ryan Passino (University of Michigan) for technical assistance with the CGNP assays. We thank members of the Department of Cell and Developmental Biology who provided access to equipment, including the O’Shea, Engel, and Spence labs. PAX6 and LIM1/2 antibodies were obtained from the Developmental Studies Hybridoma Bank, created by the Eunice Kennedy Shriver National Institute of Child Health and Human Development of the National Institutes of Health and maintained at The University of Iowa, Department of Biology, Iowa City, IA 52242, USA. Finally, we acknowledge the Biomedical Research Core Facilities Microscopy Core, which is supported by the Rogel Cancer Center, for providing access to confocal microscopy equipment.

## Author contributions

Conceptualization: B.W., B.L.A.

Data Curation: B.W., B.S.C.

Formal Analysis: B.W.

Funding Acquisition: B.W., B.L.A.

Investigation: B.W., B.S.C., B.L.A.

Methodology: B.W.

Project Administration: B.L.A.

Resources: K.J.V.

Supervision: B.L.A.

Validation: B.W.

Visualization: O.Q.M.

Writing/editing: B.W., B.L.A.

## Data and materials availability

All data are available in the main text or the supplementary materials.

**Figure S1. Timeline of *Kif17* expression during postnatal cerebellar development, validation of *Kif17* deletion, and assessment of cerebellar defects on different genetic backgrounds**.

Whole-mount X-gal stain of *Kif17*^*+/+*^ (A-C) and *Kif17*^*lacZ/lacZ*^ (D-F) cerebella from postnatal day 4 (P4) to postnatal day 21 (P21). Asterisks denote endogenous Beta Galactosidase in the choroid plexus (*1*). Scale bar (A-F), 500 μm. Fluorescent *in situ* hybridization detection of *lacZ* (yellow, H-I, K-L) in posterior cerebellar lobes of P21 *Kif17*^*+/+*^ (G-I) and *Kif17*^*-/-*^ (J-L) mice. Immunofluorescent detection of LIM1/2 (cyan) and Calbindin (CALB1, magenta) were used to visualize Purkinje cells (G, I, J, L). Scale bar, 50 μm. RT-qPCR detection of *Kif17* (M) expression in P10 *Kif17*^*+/+*^ and *Kif17*^*-/-*^ cerebella. Note that *Kif17* transcript is undetectable in *Kif17*^*-/-*^ mice. Relative expression of *Atoh1* (N) and *Kif17* (O) in P8 whole cerebella and purified cerebellar granule neural progenitor cells. Weight of cortices (P), cerebella (Q) and cerebellar weight normalized to cortical weight (R) in *Kif17*^*+/+*^, *Kif17*^*+/-*^ and *Kif17*^*-/-*^ P10 littermates on a C57BL/6J genetic background. Cerebellar weight normalized to cortical weight for *Kif17*^*+/+*^, *Kif17*^*+/-*^ and *Kif17*^*-/-*^ P10 littermates on a mixed genetic background (C57BL/6J and 129S4/SvJaeJ backgrounds, S) or a congenic 129S4/SvJaeJ genetic background (T). Data are means ± s.d. Each dot represents an individual animal or CGNP isolation. *P*-values were determined by a two-tailed Student’s *t*-test.

**Figure S2. Quantitation of cerebellar phenotypes in anterior lobes of P10 *Kif17***^***-/-***^ **mice, including reduced HH target gene expression and demonstration of reduced *Gli1* expression in P21 *Kif17***^***-/-***^ **cerebella**.

Measurements of the PCL (**A**) and EGL (**B**) thickness within anterior cerebellar lobes from P10 *Kif17*^*+/+*^ and *Kif17*^*-/-*^ mice. Percentage of Ki67^+^ (**C**) and EdU^+^ (**D**) cells in *Kif17*^*+/+*^ and *Kif17*^*-/-*^ anterior lobes in P10 cerebella. Quantitation of EdU fluorescence intensity (integrated density), within the posterior (**E**) and anterior (**F**) lobes in *Kif17*^*+/+*^ and *Kif17*^*-/-*^ P10 cerebella. Relative expression of *Ptch1* (**G**), *Ptch2* (**H**), *Ccnd1* (**I**) by RT-qPCR in P10 cerebella of *Kif17*^*+/+*^ and *Kif17*^*-/-*^ mice. Data are means ± s.d. Each dot represents the average of 3-5 images per animal (**A-F**) or an individual animal (**G-I**). *P*-values were determined by a two-tailed Student’s *t*-test. Fluorescent in situ hybridization detection of *Gli1* (yellow; **J-M**) with immunofluorescent antibody detection of PAX6 and SOX2 (magenta and cyan; **K, M**) to identify CGNs and Bergmann glia, respectively, in P21 *Kif17*^*+/+*^ (**J-K**) and *Kif17*^*-/-*^ (**L-M**) posterior cerebellar lobes. Dashed line separates Bergmann glia and CGNs. Scale bars (**K, M**), 50 μm.

**Figure S3. Validation of selective *Shh***^***Cre***^ **recombination in PCs and quantitation of cerebellar phenotypes following PC-specific *Kif17* deletion**.

Immunofluorescent analysis of *Shh*^*Cre*^-mediated recombination utilizing the *Rosa26*^*LSL-tdTomato*^ allele (**A-F**) in P10 *Rosa26*^*LSL-tdTomato*^ (**A-C**) and *Shh*^*Cre*^*;Rosa26*^*LSL-tdTomato*^ (**D-F**) cerebella. Direct fluorescence detection of tdTomato (TdTom, red; **B, C, E, F**) and antibody detection of Calbindin (CALB1, green; **A, C, D, F**) to visualize PCs. Nuclei were counterstained with DAPI (blue, **C, F**). Scale bars (**A, D**), 100 μm. Quantitation of cerebellar to cortical weights of compound *Shh/Kif17* germline mutants (**G**) and control and Purkinje cell-specific *Kif17* deletion (**H**) in P10 mice. Measurements of PCL (**I**) and EGL (**J**) thickness in anterior lobes of *Kif17*^*fl/fl*^, *Shh*^*Cre*^*;Kif17*^*+/+*^ and *Shh*^*Cre*^*;Kif17*^*fl/fl*^ P10 cerebella. Percentage of Ki67^+^ (**K**) and EdU^+^ (**L**) cells within the anterior lobes of P10 control and PC-specific *Kif17* deletion animals. Relative expression of *Ptch2* (**M**) and *Ccnd1* (**N**) by RT-qPCR in P10 whole cerebella from *Kif17*^*fl/fl*^, *Shh*^*Cre*^*;Kif17*^*+/+*^ and *Shh*^*Cre*^*;Kif17*^*fl/fl*^ mice. Data are means ± s.d. Each dot represents an individual animal (**G-H, M-N**) or average of 5 images per animal (**I-L**). *P*-values were determined by a two-tailed Student’s *t*-test.

**Figure S4. Validation of SHH antibodies, quantitation of Boc transcript/protein in *Kif17* mutant animals, and quantitation of SHH levels following *Kif17* expression in cells**.

Western blot (**A**) validation of the N-terminal SHH antibody using lysates collected from E12.5 wildtype and *Shh* mutant mouse embryos. Arrowhead indicates secreted SHH (19 kDa). Antibody detection of Calbindin (⍰ CALB1) was used to confirm equal loading across lanes. RT-qPCR analysis of *Boc* expression in *Kif17*^*+/+*^ and *Kif17*^*-/-*^ (**B**) and *Kif17*^*fl/fl*^, *Shh*^*Cre*^*;Kif17*^*+/+*^ and *Shh*^*Cre*^*;Kif17*^*fl/fl*^ (**C**) P10 cerebella. Western blot (**D**) analysis of BOC (⍰ BOC) in cerebella from adult *wildtype, Boc*^*-/-*^ and *Kif17*^*-/-*^ mice and P10 *Kif17*^*+/-*^ and *Kif17*^*-/-*^ littermates. Antibody detection of Vinculin (⍰ VIN) was used to confirm equal loading across lanes. Quantitation of the levels of BOC (**E**) normalized to Vinculin in *Kif17*^*+/-*^ and *Kif17*^*-/-*^ P10 cerebella. Quantitation of the levels of N-SHH in media (**F**) normalized to β-tubulin in COS-7 cells. Quantitation of the levels of N-SHH:GFP (**G**) and N-SHH (**H**) within COS-7 cell lysates normalized to β-tubulin. Immunofluorescence validation of C-terminal SHH antibody (green; **I-K, O-Q**) in P10 cerebella from control and *Shh* conditional *Shh* deletion animals. Antibody detection of β-galactosidase (β-GAL; red, **L-Q**) and Giantin (magenta; **O-Q**) in *Shh*^*+/+*^ (**L, O**), *Shh*^*lacZ/+*^ (**M, P**) and *Shh*^*CreER/lacZ*^ (**O, Q**) P10 posterior cerebellar lobes. Scale bar (**I-K**), 10 μm. Quantitation of SHH fluorescence (**R**) in anterior cerebellar lobes from P10 *Kif17*^*+/+*^, *Kif17*^*+/-*^ and *Kif17*^*-/-*^ mice. Data represent the mean±s.d. Individual dots represent individual mice (**B, C, E**), independent samples from a single experiment (**F-H**) or the average of 5 images per animal (**S**). *P*-values were determined by a two-tailed Student’s *t*-test.

**Figure S5. *Kif17***^***-/-***^ **CGNPs but not CGNPs from PC-specific *Kif17* deletion display increased proliferation**.

Quantitation of BrdU incorporation (**A**) in *Kif17*^*+/+*^ and *Kif17*^*-/-*^ CGNP cultures treated with either control conditioned media (control C.M.) or SHH conditioned media (SHH C.M.). Immunofluorescent analysis of *in vitro* CGNP proliferation in cultures from P8 *Kif17*^*fl/fl*^ (**B-C**) and *Shh*^*Cre*^*;Kif17*^*fl/fl*^ (**D-E**) littermates. Nuclei were counterstained with DAPI (blue); Ki67 immunofluorescence (green) and EdU incorporation (red) were visualized in response to DMSO (**B, D**) and SAG treatment (**C, E**). Scale bars (**B, D**), 100 μm. Percentage of Ki67^+^ (**F**) and EdU^+^ (**G**) cells in CGNP cultures from P8 *Kif17*^*fl/fl*^ and *Shh*^*Cre*^*;Kif17*^*fl/fl*^ littermates. Quantitation of EdU incorporation (**H**) in *wildtype, Kif17*^*-/-*^, *Boc*^*-/-*^ CGNPs grown in 10% calf serum in response to either DMSO or Smoothened Agonist (SAG, 500 nM) treatment. Each dot represents an individual well (**A, H**) or the average of 5 images per culture (**F, G**). *P*-values were determined by a two-tailed Student’s *t*-test.

**Figure S6. Validation of selective *Atoh1***^***Cre***^ **recombination in CGNPs and quantitation of cerebellar phenotypes following CGNP-specific *Kif17* deletion**.

Validation of *Atoh1Cre* specificity (**A-F**) using immunofluorescence on sections from *Rosa26*^*LSL-tdTomato*^ and *Atoh1Cre;Rosa26*^*LSL-tdTomato*^ P10 posterior cerebellar lobes. Nuclei were stained with DAPI (blue; **C, F**), and antibody detection of PAX6 (green; **B-C, E-F**) to label CGNPs and CGNs. Scale bars (**C, F**), 100 μm. P10 Cerebellar weights normalized to cortical weights (**G**) in control and CGNP-specific *Kif17* deletion animals. Measurements of PCL (**H**) and EGL (**I**) thickness in anterior lobes of *Kif17*^*fl/fl*^ and *Atoh1Cre;Kif17*^*fl/fl*^ P10 cerebella. Quantitation of the percentage of Ki67^+^ cells (**J**) and EdU^+^ cells (**K**) in the anterior lobes of P10 cerebella from *Kif17*^*fl/fl*^, and *Atoh1Cre;Kif17*^*fl/fl*^ littermates. Expression of *Ptch2* (**L**), *Ccnd1* (**M**), and *Ki67* (**N**), measured through RT-qPCR on P10 cerebella from *Kif17*^*fl/fl*^ and *Atoh1Cre;Kif17*^*fl/fl*^ littermates. Quantitation of fluorescent intensity (integrated density) of *Gli1* puncta (**O**) within CGNPs or Bergmann glia and cerebellar granule neurons (BGs + CGNs) in the anterior lobes of P10 *Kif17*^*fl/fl*^ and *Atoh1Cre;Kif17*^*fl/fl*^ mice. Each dot represents an individual animal (**G, L-M**) or the average of 5 images per animal (**H-K, O**). Data are means ± s.d. *P*-values were determined by a two-tailed Student’s *t*-test.

**Figure S7. KIF17 can physically interact with GLI transcription factors while BMP ligands attenuate CGNP proliferation**.

Relative expression of *Gli2* (**A**) and *Gli3* (**B**) measured by RT-qPCR in *Kif17*^*fl/fl*^, *Atoh1Cre;Kif17*^*+/+*^ and *Atoh1Cre;Kif17*^*fl/fl*^ in P10 whole cerebella. (**C**) Immunoprecipitation of MYC:GLI1-3 from COS-7 cells co-expressing KIF17:HA. Immunoprecipitants (IP) and whole cell lysates (WCL) were subjected to SDS-PAGE and western blot analysis (IB) using antibodies directed against MYC (α-MYC) and HA (α-HA). Antibody detection of β-tubulin (α β-TUB) was used to confirm equal loading across lanes. The molecular weights (in kDa) of protein standards are indicated at the left of each blot. Relative expression of *Atoh1* (**D**) measured by RT-qPCR in *Kif17*^*fl/fl*^, *Atoh1Cre;Kif17*^*+/+*^ and *Atoh1Cre;Kif17*^*fl/fl*^ in P10 whole cerebella. Average ciliary length of CGNPs within the posterior (**E**) and anterior (**F**) lobes of *Kif17*^*fl/fl*^ and *Atoh1Cre;Kif17*^*fl/fl*^ at P10. Relative expression of *Bmp10* (**G**) by RT-qPCR in *Kif17*^*+/+*^ and *Kif17*^*-/-*^ P10 whole cerebella. Each dot represents an individual animal (**A-B, D, G**), the average of 5 images per animal (**E-F**). Immunofluorescent analysis of *in vitro* proliferation in CGNP cultures from P8 *Kif17*^*fl/fl*^ (**H-K**) and *Atoh1Cre;Kif17*^*fl/fl*^ (**L-O**) littermates. Nuclei were counterstained with DAPI (blue); Ki67 immunofluorescence (green) and EdU incorporation (red) were visualized in response to DMSO (**H, L**), SAG (**I, M**), SAG + BMP2 (**K, N**), or SAG + BMP10 (**K, O**) treatment. Scale bars (**K, O**), 100 μm. Percentage of EdU^+^ (**R**) cells in CGNP cultures from P8 *Kif17*^*fl/fl*^ and *Atoh1Cre;Kif17*^*fl/fl*^ littermates. Ciliary lengths (**S**) measured *in vitro* from CGNP cultures from *Kif17*^*fl/fl*^ and *Atoh1Cre;Kif17*^*fl/fl*^ littermates. Each dot represents the average of 5 images per independent CGNP culture (**P**) or measurement of an individual cilium (**Q**). Data are means ± s.d. *P*-values were determined by a two-tailed Student’s *t*-test.

**Table S1. Table of antibodies used in this paper**.

**Table S2. Table of RT-qPCR primers used in this paper**.

## References

1. N. Dahmane, A. Ruiz i Altaba, Sonic hedgehog regulates the growth and patterning of the cerebellum. Development 126, 3089–3100 (1999).

2. R. J. Wechsler-Reya, M. P. Scott, Control of neuronal precursor proliferation in the cerebellum by Sonic Hedgehog. Neuron 22, 103–114 (1999).

3. P. M. Lewis, A. Gritli-Linde, R. Smeyne, A. Kottmann, A. P. McMahon, Sonic hedgehog signaling is required for expansion of granule neuron precursors and patterning of the mouse cerebellum. Dev Biol 270, 393–410 (2004).

4. J. D. Corrales, S. Blaess, E. M. Mahoney, A. L. Joyner, The level of sonic hedgehog signaling regulates the complexity of cerebellar foliation. Development 133, 1811–1821 (2006).

5. L. Izzi et al., Boc and Gas1 each form distinct Shh receptor complexes with Ptch1 and are required for Shh-mediated cell proliferation. Dev Cell 20, 788–801 (2011).

6. J. D. Corrales, G. L. Rocco, S. Blaess, Q. Guo, A. L. Joyner, Spatial pattern of sonic hedgehog signaling through Gli genes during cerebellum development. Development 131, 5581–5590 (2004).

7. F. Y. Cheng, J. T. Fleming, C. Chiang, Bergmann glial Sonic hedgehog signaling activity is required for proper cerebellar cortical expansion and architecture. Dev Biol 440, 152–166 (2018).

8. F. Bangs, K. V. Anderson, Primary Cilia and Mammalian Hedgehog Signaling. Cold Spring Harb Perspect Biol 9, (2017).

9. S. Nonaka et al., Randomization of left-right asymmetry due to loss of nodal cilia generating leftward flow of extraembryonic fluid in mice lacking KIF3B motor protein. Cell 95, 829–837 (1998).

10. S. Takeda et al., Left-right asymmetry and kinesin superfamily protein KIF3A: new insights in determination of laterality and mesoderm induction by kif3A-/-mice analysis. J Cell Biol 145, 825–836 (1999).

11. D. Huangfu et al., Hedgehog signalling in the mouse requires intraflagellar transport proteins. Nature 426, 83–87 (2003).

12. N. Spassky et al., Primary cilia are required for cerebellar development and Shh-dependent expansion of progenitor pool. Dev Biol 317, 246–259 (2008).

13. B. S. Carpenter, R. L. Barry, K. J. Verhey, B. L. Allen, The heterotrimeric kinesin-2 complex interacts with and regulates GLI protein function. J Cell Sci 128, 1034–1050 (2015).

14. N. Hirokawa, Y. Noda, Y. Tanaka, S. Niwa, Kinesin superfamily motor proteins and intracellular transport. Nat Rev Mol Cell Biol 10, 682–696 (2009).

15. M. He, S. Agbu, K. V. Anderson, Microtubule Motors Drive Hedgehog Signaling in Primary Cilia. Trends Cell Biol 27, 110–125 (2017).

16. Z. Yang, E. A. Roberts, L. S. Goldstein, Functional analysis of mouse kinesin motor Kif3C. Mol Cell Biol 21, 5306–5311 (2001).

17. X. Yin, Y. Takei, M. A. Kido, N. Hirokawa, Molecular motor KIF17 is fundamental for memory and learning via differential support of synaptic NR2A/2B levels. Neuron 70, 310–325 (2011).

18. M. F. Engelke et al., Acute Inhibition of Heterotrimeric Kinesin-2 Function Reveals Mechanisms of Intraflagellar Transport in Mammalian Cilia. Curr Biol 29, 1137–1148 e1134 (2019).

19. D. Signor, K. P. Wedaman, L. S. Rose, J. M. Scholey, Two heteromeric kinesin complexes in chemosensory neurons and sensory cilia of Caenorhabditis elegans. Mol Biol Cell 10, 345–360 (1999).

20. J. J. Snow et al., Two anterograde intraflagellar transport motors cooperate to build sensory cilia on C. elegans neurons. Nat Cell Biol 6, 1109–1113 (2004).

21. C. Insinna, N. Pathak, B. Perkins, I. Drummond, J. C. Besharse, The homodimeric kinesin, Kif17, is essential for vertebrate photoreceptor sensory outer segment development. Dev Biol 316, 160–170 (2008).

22. T. R. Lewis et al., Cos2/Kif7 and Osm-3/Kif17 regulate onset of outer segment development in zebrafish photoreceptors through distinct mechanisms. Dev Biol 425, 176–190 (2017).

23. T. R. Lewis, S. R. Kundinger, B. A. Link, C. Insinna, J. C. Besharse, Kif17 phosphorylation regulates photoreceptor outer segment turnover. BMC Cell Biol 19, 25 (2018).

24. C. Zhao, Y. Omori, K. Brodowska, P. Kovach, J. Malicki, Kinesin-2 family in vertebrate ciliogenesis. Proc Natl Acad Sci U S A 109, 2388–2393 (2012).

25. X. Yin, X. Feng, Y. Takei, N. Hirokawa, Regulation of NMDA receptor transport: a KIF17-cargo binding/releasing underlies synaptic plasticity and memory in vivo. J Neurosci 32, 5486–5499 (2012).

26. B. D. Harfe et al., Evidence for an expansion-based temporal Shh gradient in specifying vertebrate digit identities. Cell 118, 517–528 (2004).

27. E. T. Hall et al., Cytoneme delivery of Sonic Hedgehog from ligand-producing cells requires Myosin 10 and a Dispatched-BOC/CDON co-receptor complex. Elife 10, (2021).

28. A. Kawakami et al., The zebrafish-secreted matrix protein you/scube2 is implicated in long-range regulation of hedgehog signaling. Curr Biol 15, 480–488 (2005).

29. G. E. Hollway et al., Scube2 mediates Hedgehog signalling in the zebrafish embryo. Dev Biol 294, 104–118 (2006).

30. H. Y. Lee, L. A. Greene, C. A. Mason, M. C. Manzini, Isolation and culture of post-natal mouse cerebellar granule neuron progenitor cells and neurons. J Vis Exp, (2009).

31. D. Huangfu, K. V. Anderson, Cilia and Hedgehog responsiveness in the mouse. Proc Natl Acad Sci U S A 102, 11325–11330 (2005).

32. V. Matei et al., Smaller inner ear sensory epithelia in Neurog 1 null mice are related to earlier hair cell cycle exit. Dev Dyn 234, 633–650 (2005).

33. C. H. Chang et al., Atoh1 Controls Primary Cilia Formation to Allow for SHH-Triggered Granule Neuron Progenitor Proliferation. Dev Cell 48, 184–199 e185 (2019).

34. C. Cruz et al., Foxj1 regulates floor plate cilia architecture and modifies the response of cells to sonic hedgehog signalling. Development 137, 4271–4282 (2010).

35. K. A. Peterson et al., Neural-specific Sox2 input and differential Gli-binding affinity provide context and positional information in Shh-directed neural patterning. Genes Dev 26, 2802–2816 (2012).

36. Rios, R. Alvarez-Rodriguez, E. Marti, S. Pons, Bmp2 antagonizes sonic hedgehog-mediated proliferation of cerebellar granule neurones through Smad5 signalling. Development 131, 3159–3168 (2004).

37. J. Lee et al., Autoproteolysis in hedgehog protein biogenesis. Science 266, 1528–1537 (1994).

38. D. A. Bumcrot, R. Takada, A. P. McMahon, Proteolytic processing yields two secreted forms of sonic hedgehog. Mol Cell Biol 15, 2294–2303 (1995).

39. A. Porter et al., The product of hedgehog autoproteolytic cleavage active in local and long-range signalling. Nature 374, 363–366 (1995).

40. Petrov, B. M. Wierbowski, A. Salic, Sending and Receiving Hedgehog Signals. Annu Rev Cell Dev Biol 33, 145–168 (2017).

41. T. Chu, M. Chiu, E. Zhang, S. Kunes, A C-terminal motif targets Hedgehog to axons, coordinating assembly of the Drosophila eye and brain. Dev Cell 10, 635–646 (2006).

42. T. Jiwani, J. J. Kim, N. D. Rosenblum, Suppressor of fused controls cerebellum granule cell proliferation by suppressing Fgf8 and spatially regulating Gli proteins. Development 147, (2020).

43. S. Blaess, D. Stephen, A. L. Joyner, Gli3 coordinates three-dimensional patterning and growth of the tectum and cerebellum by integrating Shh and Fgf8 signaling. Development 135, 2093–2103 (2008).

44. S. Endoh-Yamagami et al., The mammalian Cos2 homolog Kif7 plays an essential role in modulating Hh signal transduction during development. Curr Biol 19, 1320–1326 (2009).

45. J. F. Dishinger et al., Ciliary entry of the kinesin-2 motor KIF17 is regulated by importin-beta2 and RanGTP. Nat Cell Biol 12, 703–710 (2010).

46. X. Wen et al., Kinetics of hedgehog-dependent full-length Gli3 accumulation in primary cilia and subsequent degradation. Mol Cell Biol 30, 1910–1922 (2010).

47. Santos, J. F. Reiter, A central region of Gli2 regulates its localization to the primary cilium and transcriptional activity. J Cell Sci 127, 1500–1510 (2014).

48. F. Haque et al., Cytoskeletal regulation of a transcription factor by DNA mimicry via coiled-coil interactions. Nat Cell Biol 24, 1088–1098 (2022).

49. Q. Ding et al., Mouse suppressor of fused is a negative regulator of sonic hedgehog signaling and alters the subcellular distribution of Gli1. Curr Biol 9, 1119–1122 (1999).

50. He et al., The kinesin-4 protein Kif7 regulates mammalian Hedgehog signalling by organizing the cilium tip compartment. Nat Cell Biol 16, 663–672 (2014).

51. M. Tuson, M. He, K. V. Anderson, Protein kinase A acts at the basal body of the primary cilium to prevent Gli2 activation and ventralization of the mouse neural tube. Development 138, 4921–4930 (2011).

52. R. Machold et al., Sonic hedgehog is required for progenitor cell maintenance in telencephalic stem cell niches. Neuron 39, 937–950 (2003).

53. J. J. Breunig et al., Primary cilia regulate hippocampal neurogenesis by mediating sonic hedgehog signaling. Proc Natl Acad Sci U S A 105, 13127–13132 (2008).

54. Y. G. Han et al., Hedgehog signaling and primary cilia are required for the formation of adult neural stem cells. Nat Neurosci 11, 277–284 (2008).

55. S. Ahn, A. L. Joyner, In vivo analysis of quiescent adult neural stem cells responding to Sonic hedgehog. Nature 437, 894–897 (2005).

56. B. Macho et al., CREM-dependent transcription in male germ cells controlled by a kinesin. Science 298, 2388–2390 (2002).

57. V. Chennathukuzhi, C. R. Morales, M. El-Alfy, N. B. Hecht, The kinesin KIF17b and RNA-binding protein TB-RBP transport specific cAMP-responsive element modulator-regulated mRNAs in male germ cells. Proc Natl Acad Sci U S A 100, 15566–15571 (2003).

58. S. Kimmins et al., A specific programme of gene transcription in male germ cells. Reprod Biomed Online 8, 496–500 (2004).

59. N. Kotaja, H. Lin, M. Parvinen, P. Sassone-Corsi, Interplay of PIWI/Argonaute protein MIWI and kinesin KIF17b in chromatoid bodies of male germ cells. J Cell Sci 119, 2819–2825 (2006).

60. M. Saade et al., Dynamic distribution of Spatial during mouse spermatogenesis and its interaction with the kinesin KIF17b. Exp Cell Res 313, 614–626 (2007).

61. A. M. Clark, K. K. Garland, L. D. Russell, Desert hedgehog (Dhh) gene is required in the mouse testis for formation of adult-type Leydig cells and normal development of peritubular cells and seminiferous tubules. Biol Reprod 63, 1825–1838 (2000).

62. Y. Wu, C. Wang, H. Sun, D. LeRoith, S. Yakar, High-efficient FLPo deleter mice in C57BL/6J background. PLoS One 4, e8054 (2009).

63. B. L. Allen et al., Overlapping roles and collective requirement for the coreceptors GAS1, CDO, and BOC in SHH pathway function. Dev Cell 20, 775–787 (2011).

64. S. Trifonov, Y. Yamashita, M. Kase, M. Maruyama, T. Sugimoto, Overview and assessment of the histochemical methods and reagents for the detection of beta-galactosidase activity in transgenic animals. Anat Sci Int 91, 56–67 (2016).

65. B. B. Madison et al., Epithelial hedgehog signals pattern the intestinal crypt-villus axis. Development 132, 279–289 (2005).

66. T. Shimokawa et al., Novel human glioma-associated oncogene 1 (GLI1) splice variants reveal distinct mechanisms in the terminal transduction of the hedgehog signal. J Biol Chem 283, 14345–14354 (2008).

67. F. Mille et al., The Shh receptor Boc promotes progression of early medulloblastoma to advanced tumors. Dev Cell 31, 34–47 (2014).

68. Y. C. Lin, S. R. Roffler, Y. T. Yan, R. B. Yang, Disruption of Scube2 Impairs Endochondral Bone Formation. J Bone Miner Res 30, 1255–1267 (2015).

69. Y. Liu et al., The BMP4-Smad signaling pathway regulates hyperandrogenism development in a female mouse model. J Biol Chem 292, 11740–11750 (2017).

70. C. H. H. Hor, J. C. W. Lo, A. L. S. Cham, W. Y. Leong, E. L. K. Goh, Multifaceted Functions of Rab23 on Primary Cilium-Mediated and Hedgehog Signaling-Mediated Cerebellar Granule Cell Proliferation. J Neurosci 41, 6850–6863 (2021).

71. M. K. Scales et al., Combinatorial Gli activity directs immune infiltration and tumor growth in pancreatic cancer. PLoS Genet 18, e1010315 (2022).

72. Y. Han et al., Regulation of Gli ciliary localization and Hedgehog signaling by the PY-NLS/karyopherin-beta2 nuclear import system. PLoS Biol 15, e2002063 (2017).

