## Supplementary figures and images for "Dual and Opposing Roles for the Kinesin-2 Motor, KIF17, in Hedgehog-dependent Cerebellar Development"

### Supplemental Figure 1

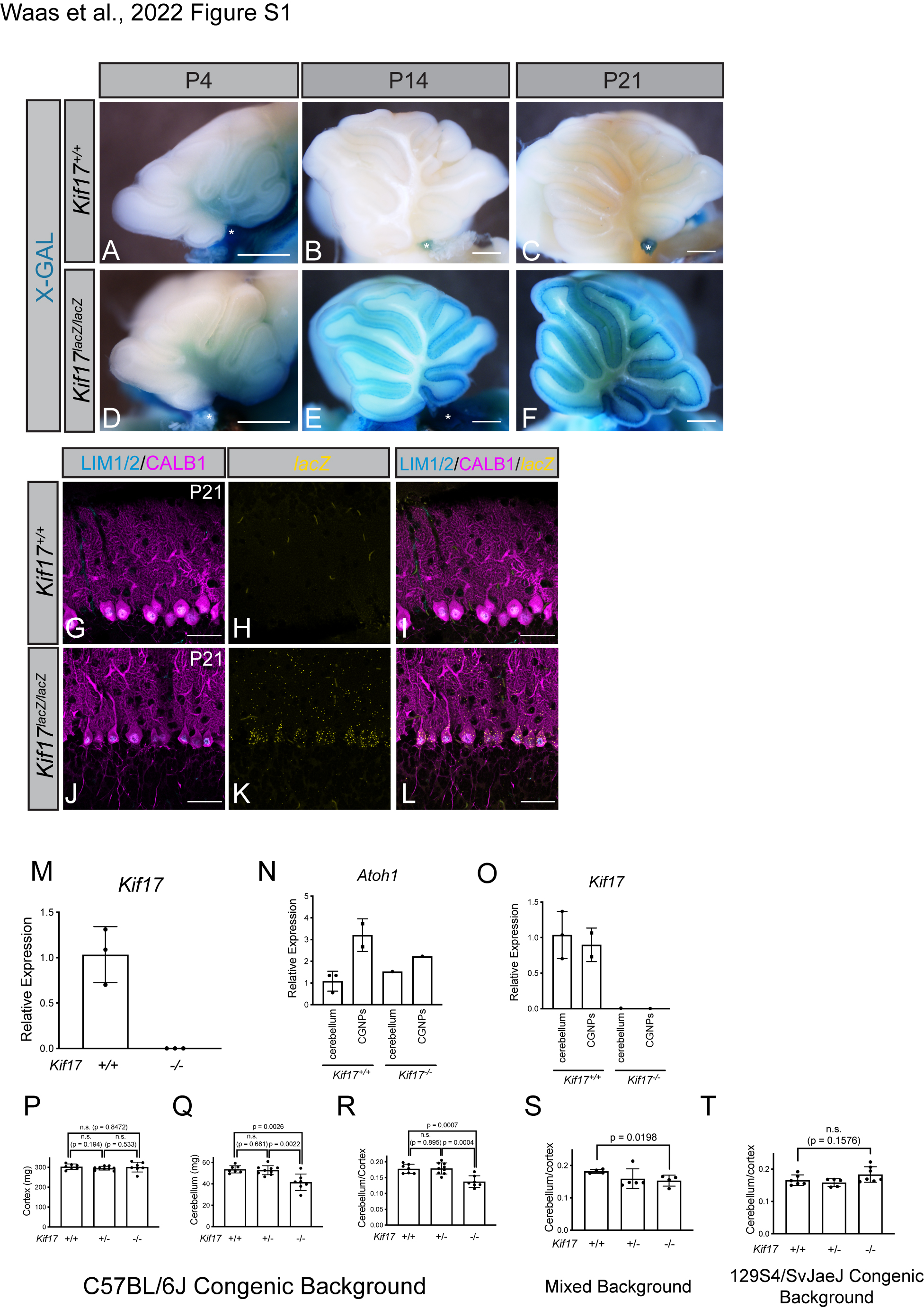

### Supplemental Figure 2

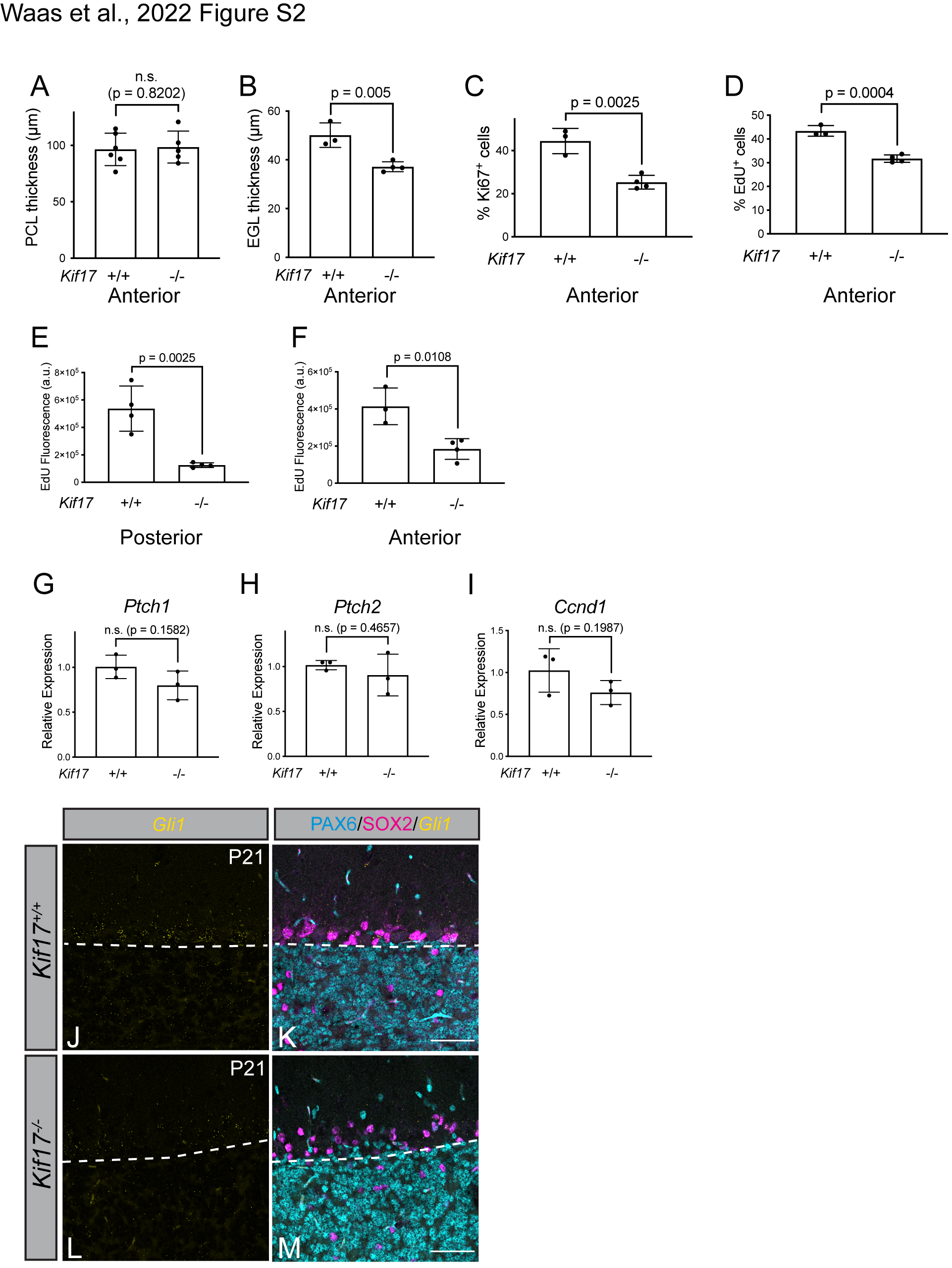

### Supplemental Figure 3

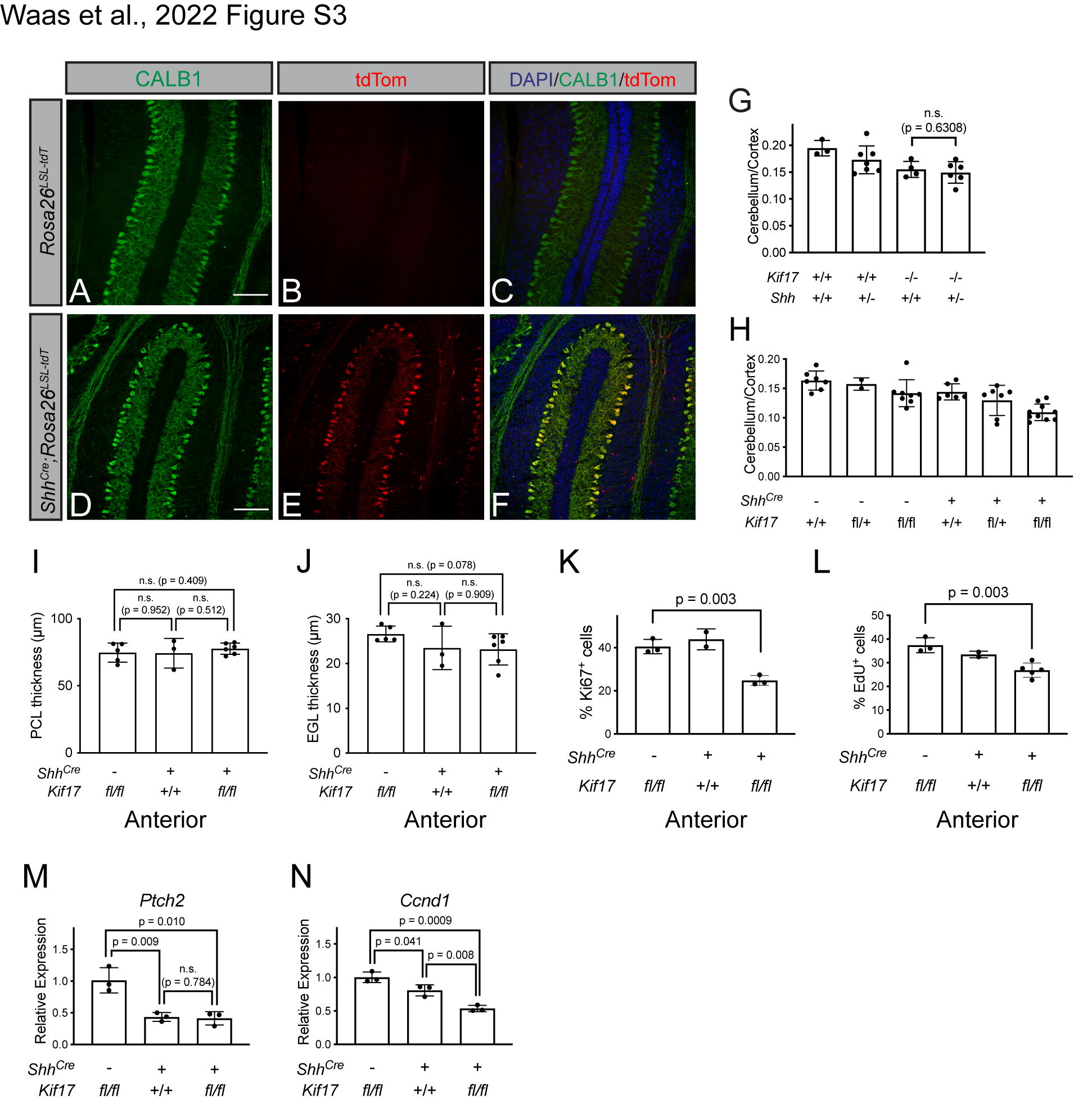

### Supplemental Figure 4

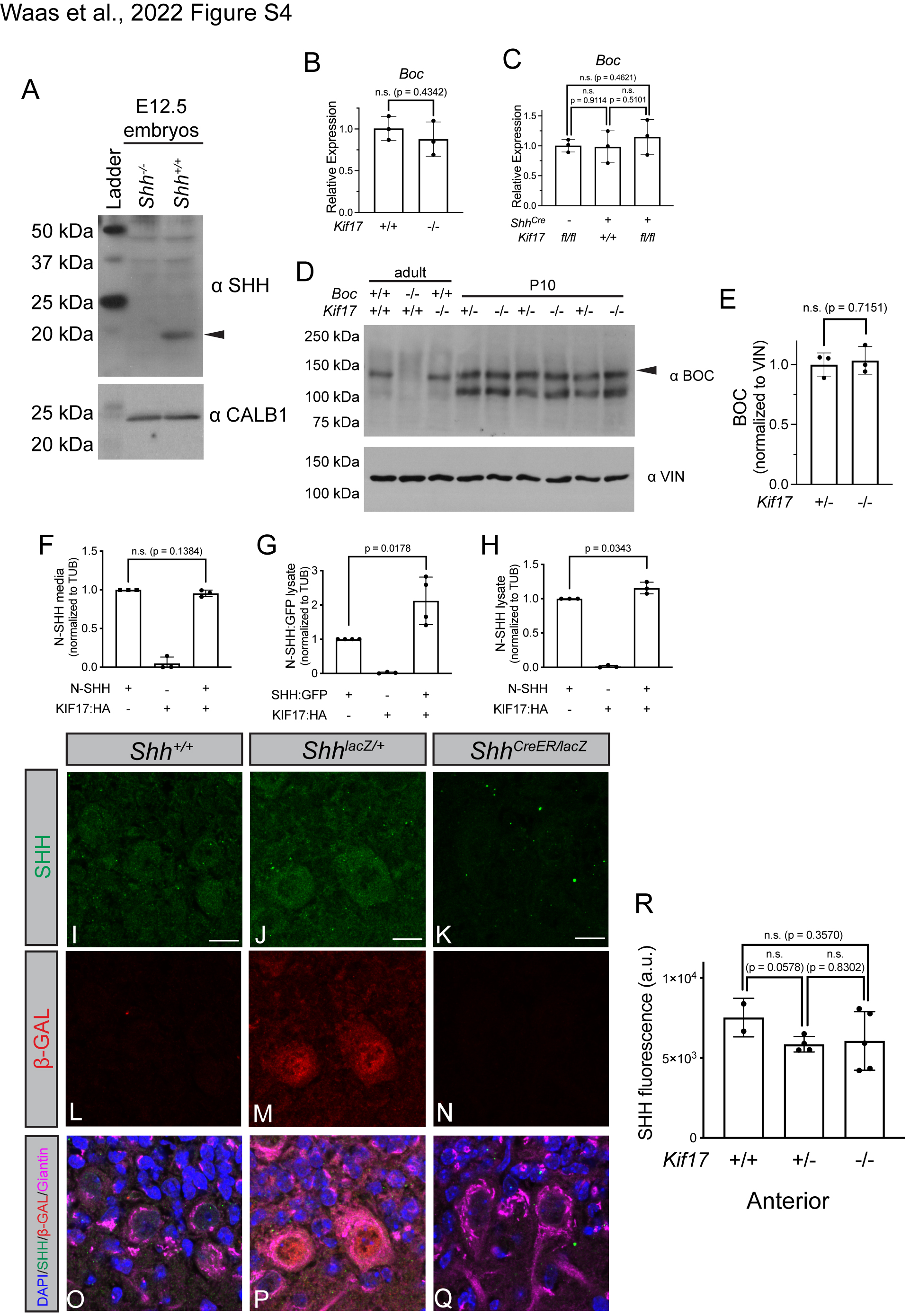

### Supplemental Figure 5

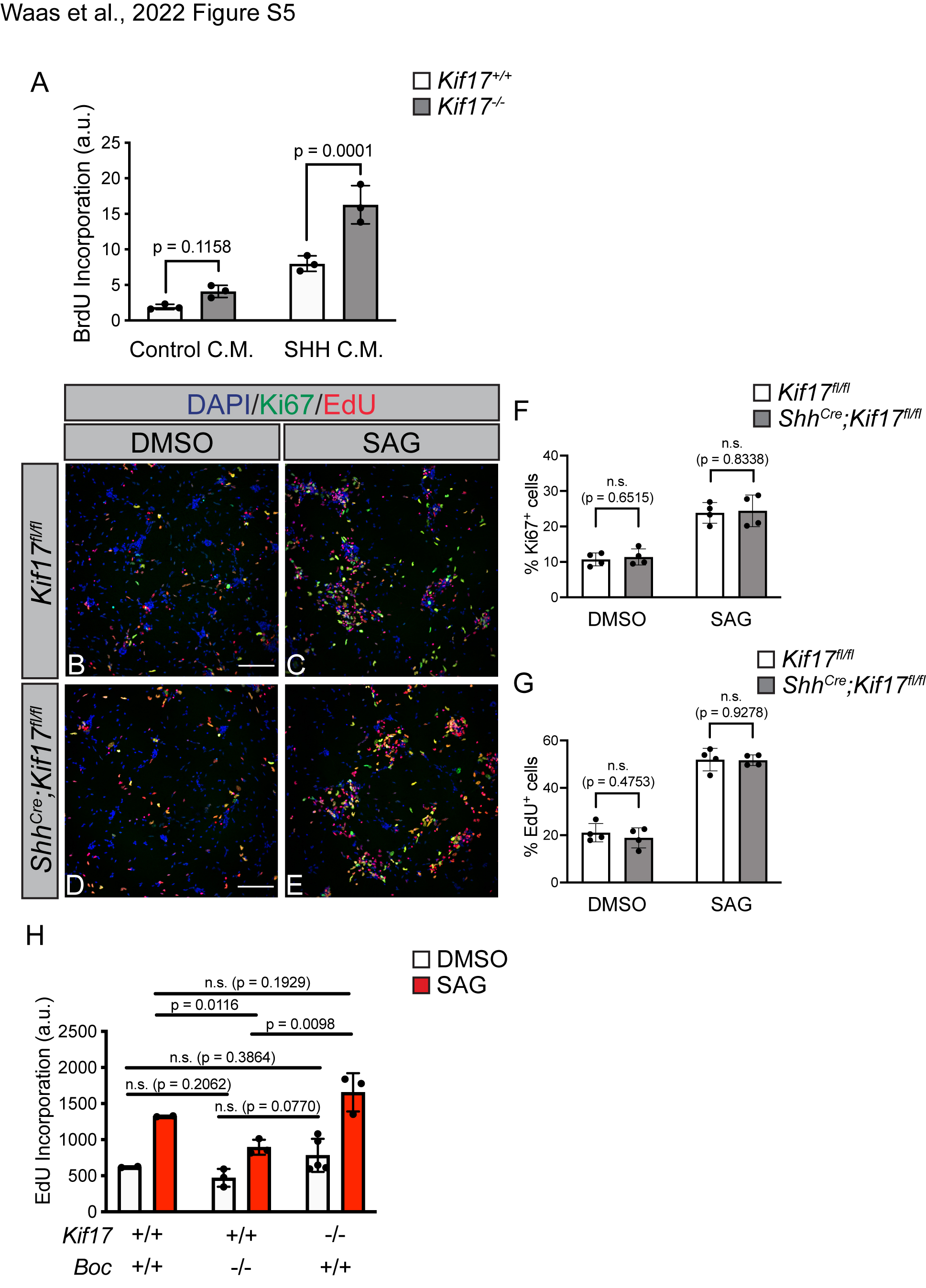

### Supplemental Figure 6

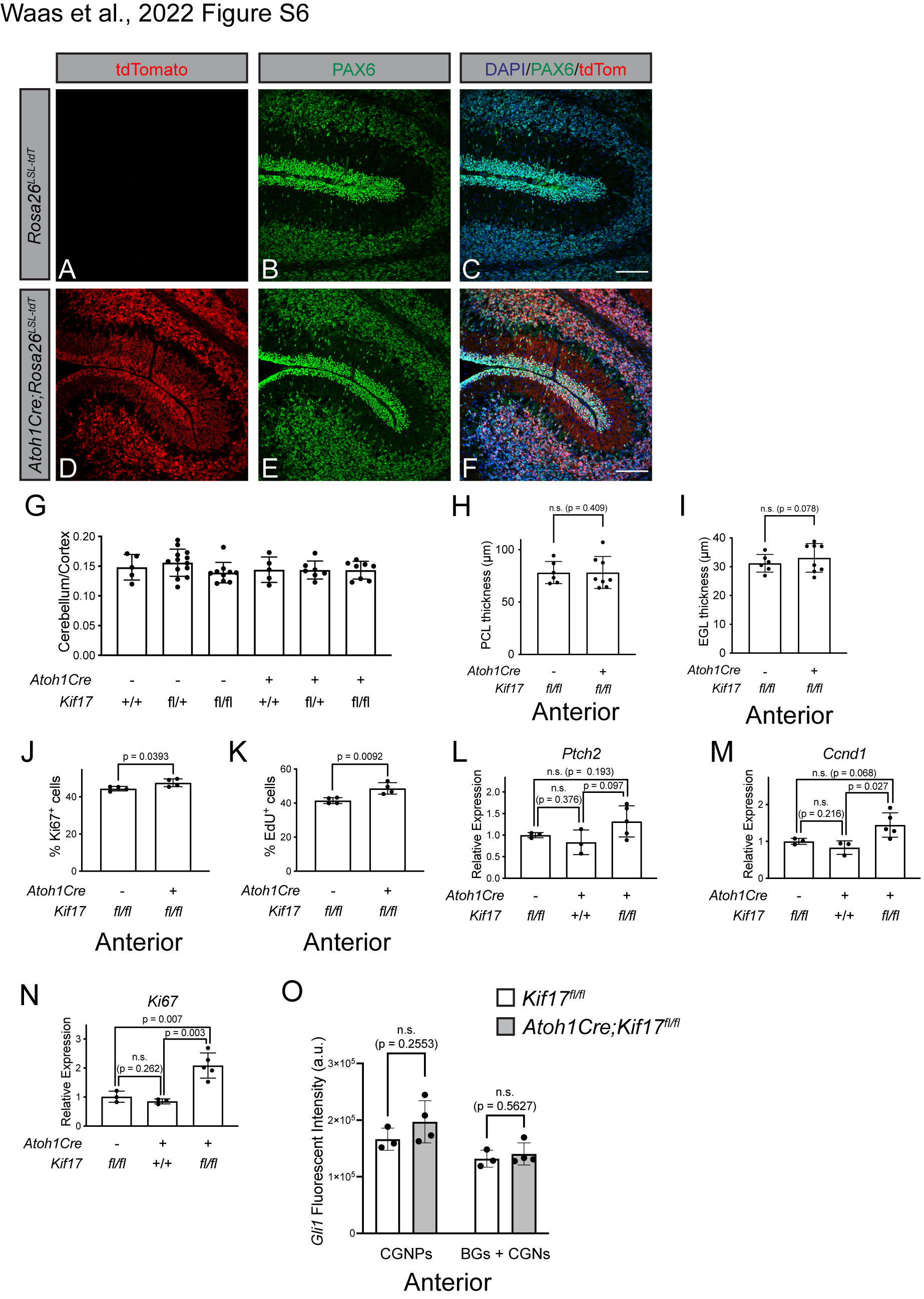

### Supplemental Figure 7

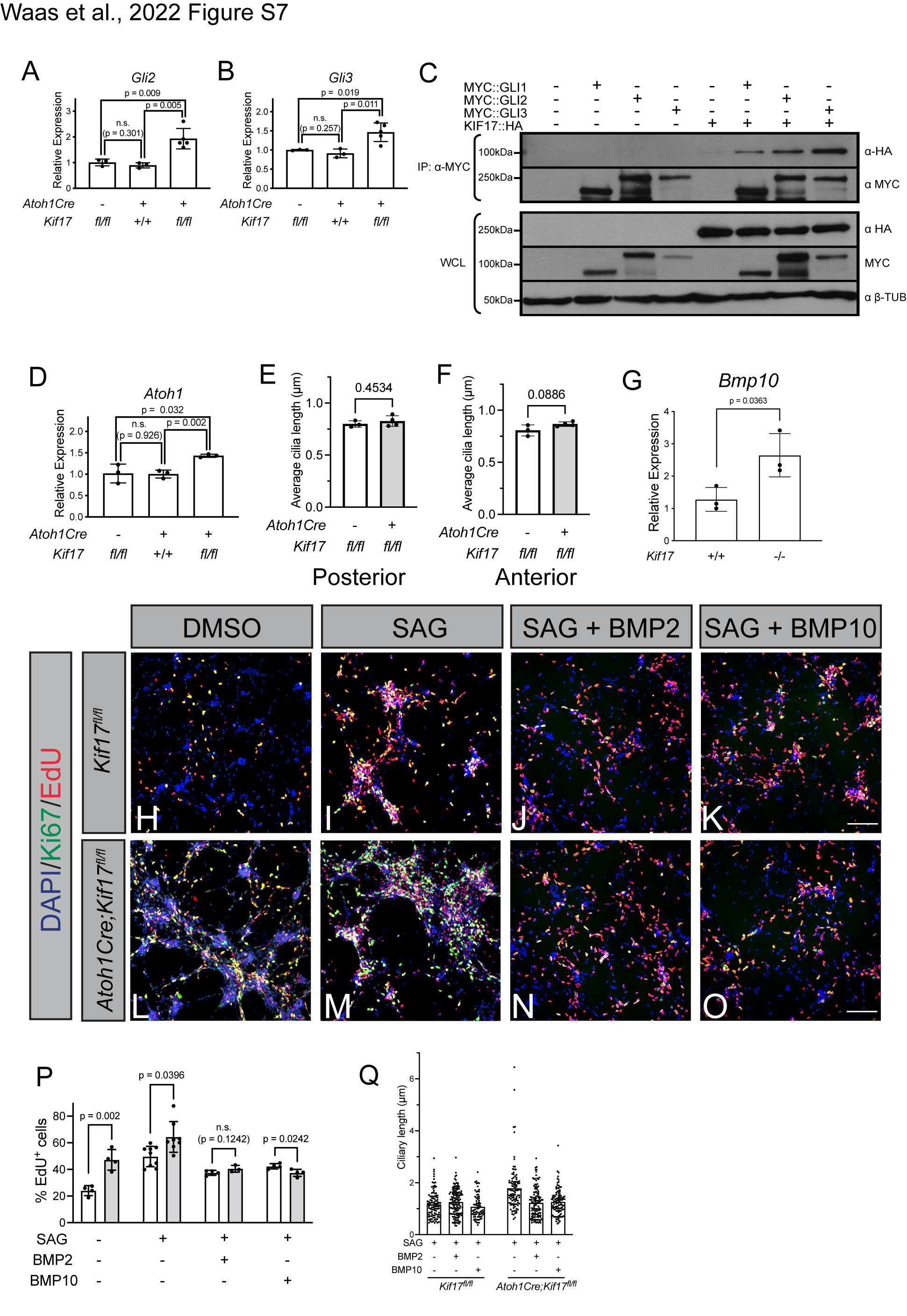
