## Supplemental Table 1 for "Dual and Opposing Roles for the Kinesin-2 Motor, KIF17, in Hedgehog-dependent Cerebellar Development"

| Antibody | Source | Catalogue Number | Application | Concentration used |
|---|---|---|---|---|
| Mouse IgG1 anti PAX6 | DSHB | PAX6 | IF | 1:20 on tissue sections, 1:40 on coverslips |
| Rabbit anti Calbindin (CALB-1) | SWANT | CB38 | IF/WB | 1:10,000 (IF), 1:2000 (WB) |
| Chicken anti Beta-galactocidase | ICL | CGAL-45A-Z | IF | 1:2000 |
| Mouse IgG1 anti LIM1+2 | DSHB | 4F2 | IF | 1:20 |
| Rabbit anti Ki67 | Abcam | ab15580 | IF | 1:1000 on tissue sections, 1:2000 on coverslips |
| Rabbit anti SOX2 | Seven Hills Bioreagent | WRAB-1236 | IF | 1:2000 |
| Goat anti SHH (N-terminus) | R&D systems | AF464 | WB | 0.5 $\mu$g/mL |
| Rabbit anti HA | Bethyl Labs | A190-108A | WB | 1:10,000 |
| Mouse IgG1 anti Beta-tubulin (B-Tub) | DSHB | E7 | WB | 1:2000 |
| Goat anti SHH (C-terminus) | R&D systems | AF445 | IF | 10 $\mu$g/mL |
| Goat anti BOC | R&D systems | AF2385 | WB | 1:4000 |
| Rabbit anti Giantin | Biolegend (Covance) | 924302 | IF | 1:1000 |
| Goat anti GLI3 | R&D systems | AF3690 | WB | 1:1000 |
| Rabbit anti GLI1 | Cell Signaling Technology | 2534 | WB | 1:1000 |
| Goat anti GLI2 | R&D systems | AF3635 | WB | 1:1000 |
| Rabbit anti VINCULIN | Cell Signaling Technology | 13901 | WB | 1:1000 |
| Mouse IgG1 anti MYC | Santa Cruz | sc-40 | IP/WB | 1:150 (IP), 1:1000 (WB) |
| Mouse IgG1 anti HA | Covance | MMS-101 | IP/WB | 1:300 (IP), 1:1000 (WB) |
| Mouse IgG2a anti ARL13B | NeuroMAB | 73-287 | IF | 1:100 on tissue sections, 1:200 on coverslips |

| | | | | |
|---|---|---|---|---|
| Rabbit anti Gamma-Tubulin | Sigma | T3559 | IF | 1:4000 on tissue sections, 1:8000 on coverslips |
| Alexa Fluor 488 goat anti-mouse IgG1 | Invitrogen | A21121 | IF | 1:500 |
| Alexa Fluor 647 goat anti-mouse IgG1 | Invitrogen | A21240 | IF | 1:500 |
| Alexa Fluor 647 donkey anti-rabbit IgG | Invitrogen | A31573 | IF | 1:500 |
| Cy3 AffiniPure Donkey anti-Chicken IgY | Jackson Immunoresearch | 703-165-155 | IF | 1:500 |
| AP-conjugated anti-DIG antibody | Roche (Millipore Sigma) | 11093274910 | SISH | 1:4000 |
| Polyclonal Donkey anti Goat HRP | R&D systems | HAF109 | WB | 1:1000-5000 |
| Peroxidase AffiniPure Donkey Anti-Rabbit IgG | Jackson Immunoresearch | 711-035-152 | WB | 1:5000 |
| Peroxidase AffiniPure Donkey Anti-Mouse IgG | Jackson Immunoresearch | 715-035-150 | WB | 1:5000 |
| Alexa Fluor 488 donkey anti-goat IgG | Invitrogen | A11055 | IF | 1:500 |
| AffiniPure goat anti-mouse-light-chain secondary antibody | Jackson Immunoresearch | 115-035-174 | WB | 1:50,000 |
| Alexa Fluor 488 goat anti-mouse IgG2a | Invitrogen | A21131 | IF | 1:500 |
