## Supplemental Table 2 for "Dual and Opposing Roles for the Kinesin-2 Motor, KIF17, in Hedgehog-dependent Cerebellar Development"

| RT-qPCR Primers | | | |
| --- | --- | --- | --- |
| Gene | forward primer (5-3) | reverse primer (5-3) | Source |
| *Gapdh* | GTGGTGAAGCAGGCATCTGA | GCCATGTAGGCCATGAGGTC | Han et al., 2017 (PLoS Biology) |
| *Kif17* | CATGCACACGGTACACAAC | GAACGGGAGGAGTCCTTATTC | designed by BW |
| *Atoh1* | AGTCAATGAAGTTGTTTCCC | ACAGATACTCTTATCTGCCC | Hor et al., 2021 (Journal of Neuroscience) |
| *Gli1* | GTGCACGTTTGAAGGCTGTC | GAGTGGGTCCGATTCTGGTG | Han et al., 2017 (PLoS Biology) |
| *Ptch1* | gaagccacagaaaaccctgtc | gccgcaagccttctctagg | Han et al., 2017 (PLoS Biology) |
| *Ptch2* | CCCGTGGTAATCCTCGTGGCCTCTAT | TCCATCAGTCACAGGGGCAAAGGTC | Shimokawa et al., 2008 (JBC) |
| *Ccnd1* | AGACCTGTGCGCCCTCCGTA | CAGCTGCAGGCGGCTCTTCT | Han et al., 2017 (PLoS Biology) |
| *Shh* | CATGGTCTCGCTGCTCAA | CAGCTGCTCTACCACATTGGCACC | |
| *Scube2* | TGACTACCTGGTGATGCGGAAAAC | CAGTGGCGTGTGGGAAGAGTCA | Lin et al., 2015 (J Bone Miner Res.) |
| *Boc* | TTCATCCCCTTCTGCCTATG | ACCATTGTGTACTGGCACGA | Mille et al., 2014 (Dev Cell) |
| *Scube1* | CGGCGGCGAACTTGGTGACTACA | TTGATAAAGGACCGGGGGAACAT | Lin et al., 2015 (J Bone Miner Res.) |
| *Scube3* | TGCTCCCCGGGCCACTACTAT | AGCGCTGTTGGCCTCACTGGTCTT | Lin et al., 2015 (J Bone Miner Res.) |
| *Ki67* | CATTGACCGCTCCTTTAGGTATGAAG | TTGGTATCTTGACCTTCCCCATCAG | Mille et al., 2014 (Dev Cell) |
| *Gli2* | CCTTCACCCACCTTCTTGG | CTTGTTCTGGTTGGCATCATTT | Scales et al., 2022 (in press) |
| *Gli3* | CACATGCATCAACAGATCCTAAGC | AGGGATAGGTCTCTGTGTTGGAAAT | Scales et al., 2022 (in press) |
| *Bmp2* | GGGACCCGCTGTCTTCTAGT | TCAACTCAAATTCGCTGAGGAC | Liu et al., 2017 (J Biol Chem.) |
| *Bmp4* | TTCCTGGTAACCGAATGCTGA | CCTGAATCTCGGCGACTTTTT | Liu et al., 2017 (J Biol Chem.) |
| *Bmp10* | ATGGGGTCTCTGGTTCTGC | CAATACCATCTTGCTCCGTGAA | Liu et al., 2017 (J Biol Chem.) |
